# Hippocampal Neuroinflammation and Altered Peripheral Neurobiological Protein Profile in Experimental Arthritis and Systemic Juvenile Idiopathic Arthritis

**DOI:** 10.64898/2026.03.13.711607

**Authors:** Xingzhao Wen, Heshuang Qu, Malgorzata Benedyk-Machaczka, Daphne Chen, Erik Sundberg, Erik Melén, Maria Altman, Cecilia Aulin, Helena Erlandsson Harris

## Abstract

**Background:** Children with juvenile idiopathic arthritis (JIA) are reported to exhibit increased rates of symptoms affecting emotional regulation and behavior. However, underlying biological mechanisms remain unclear. Neuroinflammation in the central nervous system (CNS) can be triggered by peripheral immune effects and may contribute to these observations. In this study, we aimed to investigate if neurobiological alterations are present in systemic JIA (sJIA), and if CNS neuroinflammation occurs during arthritis, and to explore the potential mechanisms involved.

**Methods:** Plasma samples from patients with active sJIA (n = 16) and sex- and age-matched healthy controls (HCs, n = 16), together with paired samples from the same sJIA patients during inactive disease (n = 12), were analyzed using Olink proteomics to determine the peripheral neurobiological and inflammation protein profiles. Clinical data was retrieved from the Swedish Pediatric Rheumatology Register and medical charts. CNS Neuroinflammatory responses and underlying mechanisms were further explored through *in vivo* and *in vitro* experiments.

**Findings:** Active sJIA patients exhibited altered neurobiological protein profiles compared with HCs. These alterations correlated with higher scores of pain and life impact in patients, suggesting that the altered profiles may reflect neurofunctional changes in the patients. Notably, the neurobiological protein profile remained altered even during the inactive phase of the disease. In chronic arthritic mice, microglial activation and impaired neurogenesis were observed in hippocampus, with no significant cortical changes. RNA-seq analysis implicated mitochondrial dysfunction and oxidative stress in mediating neuroinflammation during chronic arthritis in mice. Heme oxygenase 2 (HMOX2) was identified as a peripheral biomarker indicating hippocampal microglia activation. Combined neurobiological and inflammation profiling in sJIA patients implicated Interleukin-6 (IL-6) and Interleukin-18 (IL-18) as key drivers of hippocampal microglia activation during arthritis.

**Interpretation:** Chronic arthritis is associated with neuroinflammation and altered neurobiological protein profiles in sJIA. HMOX2 emerges as a promising plasma biomarker of CNS changes. IL-6 and especially IL-18 are indicated as key drivers of neuroinflammatory processes. These findings offer insights for clinical monitoring and targeted therapies.

**Funding:** This study was funded by grants from the Swedish Research Council and The Swedish Rheumatism Association.

**Research in context:** *Evidence before this study:* Children with juvenile idiopathic arthritis (JIA) have increased rates of emotional and behavioral disturbances compared with healthy peers. Systemic inflammation and chronic arthritis are suspected to affect the central nervous system, but biological mechanisms in systemic JIA (sJIA) are poorly understood.

*Added value of this study:* In this study, we demonstrate patients with sJIA have a distinct plasma neurobiological protein profile compared with healthy controls, which correlate with higher pain and life impact scores. In chronic arthritic mice, hippocampal microglial activation, impaired neurogenesis, and mitochondrial dysfunction with oxidative stress are presented. By combining patient and mouse data, we identify heme oxygenase 2 (HMOX2) as a candidate plasma biomarker of hippocampal neuroinflammation and implicate IL-6, and especially IL-18, as key mediators linking chronic arthritis to neurobiological changes.

*Implications of all the available evidence:* This study provides molecular evidence that neurobiological alterations in sJIA patients and supports incorporating neurobiological and neuropsychiatric monitoring into the clinical follow-up of children with sJIA. We highlight the mechanistic targets and measurable biomarkers (e.g. HMOX2) for future studies and trials aiming to modulate neuroinflammation in chronic arthritis. This study may inform the development of personalized treatment strategies, including IL-18–directed therapies, for patients at risk of neurological or psychosocial complications.

## 1. BACKGROUND

Juvenile idiopathic arthritis (JIA) is the most common chronic rheumatic disease in children under 16, characterized by persistent joint inflammation and significant impact on quality of life [1, 2]. It comprises seven subtypes defined by the International League of Associations for Rheumatology (ILAR) [3]. Although the clinical features vary among JIA subtypes, all patients exhibit inflammation, joint destruction, pain, and fatigue [3]. The subtype systemic JIA (sJIA) is the most severe form of JIA, with flares of systemic inflammation in addition to arthritis [4].

Beyond musculoskeletal symptoms, clinicians have also observed psychiatric impairment in patients with JIA[5–9]. Children with JIA have been reported to exhibit higher rates of depression and anxiety compared to their healthy peers[5, 8, 9]. Lower scores in neurocognitive performance tests were reported in JIA patients as compared with age-matched healthy individuals [7]. In addition, as adults, these patients continue to display cognitive differences relative to the general population, which can significantly impact their work and daily life and may persist throughout their lifetime[10, 11]. For JIA patients, psychiatric and cognitive impairments are commonly associated to factors like chronic pain, social limitations and restricted physical activities[6, 8, 12]. Recently, a few studies have suggested that neuroinflammation in the CNS may occur in patients with JIA and could contribute to psychiatric dysfunction[13, 14]. Despite these observations, the pathological changes in the CNS associated with JIA remain poorly understood. Furthermore, the mechanisms underlying CNS involvement in JIA remain largely unknown. Currently, there are no specific treatment or biomarkers available for neuropsychiatric manifestations in JIA.

Neuroinflammation, inflammation in the CNS, has been identified as a contributing factor to cognitive decline and psychiatric disorders[15, 16]. Neuroinflammation has been suggested to precipitate alterations in neurotransmitter systems, leading to behavioral and cognitive changes [17]. During neuroinflammation, resident CNS cells such as microglia, astrocytes, and oligodendrocytes become activated and contribute to the inflammatory response by releasing cytokines, chemokines, damage-associated molecular patterns (DAMPs), and reactive oxygen species (ROS), which in turn can damage brain tissue and influence the onset of psychiatric manifestations [18, 19]. More importantly, increased evidence suggests that neuroinflammation can be triggered by peripheral effectors, such as autoimmune diseases, infection, and surgery [20, 21].

In this study, we collected plasma samples from sJIA patients to investigate whether the profile of neurobiological related proteins is altered compared with that of healthy children. In parallel, we used a chronic arthritis mouse model to explore if CNS neuroinflammation presents during chronic arthritis and to investigate underlying mechanisms linking arthritis and neuroinflammation.

## 2. MATERIALS AND METHODS

### 2.1 Ethics approval

Samples from patients with JIA and from healthy children were collected in accordance with the Declaration of Helsinki. Ethical approval was obtained from the North Ethical Committee in Stockholm, Sweden (Dnrs 2009-1139-30-4 and 2010-165-31-2 for JIA patient samples; Dnr 03-067 for the healthy controls). All animal procedures were performed in compliance with protocols approved by the First Regional Ethical Committee for Animal Experiments in Kraków, Poland (approval number: 329/2022).

### 2.2 Clinical study population

Plasma samples were collected from 16 patients with active sJIA at Astrid Lindgren’s Children’s Hospital, Karolinska University Hospital, Stockholm, Sweden (sample collection JABBA). 16 age- and sex-matched healthy individuals (Barnens Miljö- och Hälsoundersökning cohort) from the Stockholm region were selected and were matched to patients on a one-to-one basis by sex and age category. Because the healthy control cohort primarily included children aged 4, 8, and 12 years, exact age matching was not feasible for all patients. Therefore, sJIA patients were grouped into predefined age bands (1–5 years, 6–10 years, and ≥11 years), and each patient was matched to a sex-matched control within the corresponding age category.

For 12 of the 16 sJIA patients sampled during active disease, samples from later inactive disease states were available and were also collected for comparative analysis. Clinical and laboratory data collected at each sampling occasion were retrieved from medical records and from the Swedish Pediatric Rheumatology Quality Register (Svenska Barnreumaregistret, Omda®) and are presented in Table S1.

### 2.3 Proteomics assays

Plasma samples were subjected to proteomic profiling using the Olink Target 96 Neuro-exploratory Panel (Olink Bioscience, Sweden), high-throughput multiplex immunoassays based on proximity extension assay (PEA) technology. The Olink Target 96 Neuro-Exploratory panel measures 92 proteins involved in diverse neurobiological processes and disease mechanisms (https://olink.com/products/olink-target-96). We also mapped inflammatory proteins profile using the Inflammation panel includes 92 proteins associated with inflammatory and immune response pathways[22]. Protein abundance was quantified as log2-normalized protein expression (NPX) values. Proteins with NPX values below the limit of detection (LOD) in more than 80% of the samples were excluded. Differentially expressed proteins (DEPs) between groups were defined as those with a ΔNPX > 1 (fold change > 2) and an adjusted P value (Padj) < 0.05. Adjusted P values were calculated using the Benjamini–Hochberg procedure to control the false discovery rate (FDR) at 5% within each comparison.

Principal component analysis (PCA) was performed on normalized protein expression data. The first three principal components were visualized in a three-dimensional PCA plot to assess sample clustering and overall variance structure. Hierarchical clustering analysis was performed using the ClustVis web tool (https://biit.cs.ut.ee/clustvis/) with Euclidean distance on z-score–normalized protein expression values from the Neuro Exploratory panel, and clustering was conducted using Ward’s linkage.

### 2.4 Induction of experimental arthritis and tissue collection

Female DBA/1 mice (8–10 weeks old) were obtained from Janvier Labs (France), and were housed at the Jagiellonian University in Krakow, Poland. Mice were randomly assigned to experimental groups using a random number method prior to model induction. Collagen-induced arthritis (CIA) was induced in female DBA/1 mice as previously described[23]. Briefly, on day 0, 10-week-old mice were immunized with 100□µl of bovine type II collagen (CII; 100□µg, Chondrex, Inc., USA) emulsified in complete Freund’s adjuvant (CFA; 50□µg Mycobacterium tuberculosis, Chondrex, Inc., USA) via intradermal injection at the base of the tail. 4 weeks later, a booster injection of 100□µl of CII (100□µg) emulsified in incomplete Freund’s adjuvant (IFA, Chondrex, Inc., USA) was administered. Control mice received intradermal phosphate-buffered saline (PBS) injections at day 0 and 4 weeks later. Mice were monitored three times per week for signs of joint inflammation and scored using a 12-point scale, with each paw scored from 0 to 3 based on swelling and redness (0 = normal, 1 = mild, 2 = moderate, 3 = severe), giving a maximum possible score of 12 per mouse [24]. The study was terminated on day 90.

Upon termination, mice were deeply anesthetized with isoflurane (Baxter, UK) and sacrificed by transcardial perfusion with cold PBS. Before perfusion, Blood was collected via cardiac puncture and allowed to clot at room temperature for 30 minutes. Samples were then centrifuged at 2000 × g for 10 minutes at 4□°C, and the serum was aliquoted and stored at −80□°C.

After perfusion, Brain tissues were bisected along the midline. One half was used to isolate the hippocampus and cortex for subsequent RNA and protein extraction. The other half of the brain tissues were immersed in 4% paraformaldehyde (PFA, Thermo Fisher, USA) in PBS for 24 hours at room temperature. Then, tissues were cryoprotected in 20% sucrose in PBS for 48 hours at 4°C. Tissues were embedded in optimal cutting temperature compound (OCT, Leica Biosystems, USA) and then stored at -80°C freezer until sectioning. For the sectioning, tissues were cut into slices of 16 µm or 8µm with the Cryostar NX7 (Thermo Fisher, USA). Slices were mounted on SuperFrost Plus adhesion glass slides (Thermo Fisher, USA) and stored at -80°C for staining analysis.

### 2.5 RNA and protein extraction from tissues

Hippocampus tissues were dissected and used for further RNA and protein extraction. Tissues were homogenized with beads in 1ml Qiazol lysis reagent (Qiagen, Germany) using the TissueLyser II (Qiagen, Germany). 200μl of chloroform was added, followed by vigorous shaking for 15 seconds. The mixture was incubated at room temperature for three minutes and centrifuged at 12,000 × g for 15 minutes at 4□°C to separate the phases. The aqueous phase was carefully transferred to a new tube for RNA precipitation. The interphase and organic phase were retained for protein extraction.

For RNA extraction, the RNeasy Lipid Tissue Kit (Qiagen, Germany) was used to isolate RNA from the tissues, according to manufacturer’s protocol. The purity and concentration of RNA were measured by QIAxpert (Qiagen, Germany).

For protein extraction, 300 μl of 100% ethanol (Solveco□AB, Sweden) was added to the interphase/organic phase, mixed thoroughly, and incubated for 3 minutes at room temperature, followed by centrifugation at 2,000 × g for 5 minutes at 4□°C. The supernatant was transferred to a new tube, and proteins were precipitated by adding 1.5 ml of isopropanol (Solveco□AB, Sweden). After incubation for 10 minutes at room temperature, the samples were centrifuged at 12,000 × g for 10 minutes at 4□°C. The resulting protein pellet was washed three times with 0.3 M guanidine hydrochloride (Sigma-Aldrich, USA) in 95% ethanol, followed by a final wash with 100% ethanol. The pellet was air-dried and resuspended in 1% sodium dodecyl sulfate (SDS) buffer for western blotting analysis.

### 2.6 RNA sequencing and data analysis

RNA samples originating from 6 arthritic mice and 6 healthy controls were used for RNA sequencing analysis. Sequencing and library preparation were carried out at the Genomic Core Facility at the University of Bergen. Library preparation was performed using Illumina stranded mRNA preparation ligation kit with a total of 200 ng RNA. Illumina 10 bp unique dual indexes (UDI) were used and a total of eleven polymerase chain reaction (PCR) cycles were run. Briefly, mRNA was enriched from total RNA using poly(A) selection and converted into strand-specific libraries following the manufacturer’s instructions. Sequencing was performed on an Illumina NovaSeq 6000 platform using a NovaSeq SP flowcell with a paired-end 2 × 100 bp setup. Raw data were processed using the standard Illumina pipeline to produce demultiplexed FASTQ files for downstream analysis. FASTQ files were subjected to quality control using FastQC. The results from all samples were aggregated using MultiQC (v1.11) to assess overall data quality. Clean reads were then aligned to the mouse genome (GRCm38.p5) using HISAT2 (v2.0.5) with the Gencode vM13 gene transfer format (GTF) file as the reference annotation. Gene-level counts were obtained using FeatureCounts (v1.5.2). The count matrix was generated and was used for downstream analysis.

Downstream analyses were performed using R software (v4.4.1). Briefly, genes with zero counts across all samples and non-protein coding genes were excluded from the data analysis. Normalization was conducted using the median-of-ratios method implemented in DESeq2, with group (arthritic vs. healthy control mice) specified in the design formula (∼ group). Differential expression was assessed using the Wald test. Raw p-values were adjusted for multiple testing using the Benjamini–Hochberg procedure as implemented in DESeq2 to control the FDR. Gene identifiers were further annotated with gene symbols and descriptions using Biomart (v2.61.3). To capture potentially relevant biological signals, genes were considered differentially expressed when adjusted p-value (FDR) was < 0.1 and the log2 fold change was < - 0.585 or > 0.585.

Three-dimensional PCA plots were generated using the plotly package (v4.10.3) in R, based on the complete RNA-seq expression matrix. Gene set enrichment analysis (GSEA) was performed using the mouse Molecular Signature Database (MSigDB) [25], applying the Gene Ontology biological process (GO:BP) collection to a pre-ranked list of all genes from the RNA-seq dataset. Analysis was performed and results were visualized with GSEA software, version 4.3.3.

### 2.7 Immunofluorescence staining

Before staining, the OCT-embedded tissue sections were dried at room temperature for 15 min. Antigen retrieval was performed using a sodium citrate buffer (sodium citrate 10 mM, 0.05% Tween-20, pH=6.0), with one retrieval cycle of the 2100 antigen retriever (Aptum Biologics, UK). Sections were blocked in 0.3% Triton X- 100 (Sigma-Aldrich, USA) and 5% serum (the type of serum depended on the antibodies used). Sections were incubated with the primary antibodies at 4 ℃ overnight. Next, sections were washed with PBS containing 0.02% Tween-20 before incubation with a secondary antibody for 1-2 hours at room temperature. After another wash, sections were incubated with 0.1% Sudan Black B (Sigma-Aldrich, USA) in 70% ethanol for 20 minutes and room temperature. Next, slides were mounted using ProLong™ Gold Antifade Mountant with DNA Stain DAPI (Thermo Fisher, USA). Images were captured with the Zeiss LSM-880 confocal microscope. Fluorescence intensity quantification was performed using ImageJ (version 1.54). The details of primary and secondary antibodies are listed in Table S2.

### 2.8 Image analysis

Microglial morphology was analyzed using the Skeleton analysis plugins in Fiji/ImageJ software (NIH, USA). Briefly, confocal z-stack images of Iba1-stained microglia were first converted to 8-bit grayscale, and brightness/contrast was adjusted to optimize visualization. Images were then processed with a Gaussian blur filter to enhance continuous structures, followed by thresholding to generate binary masks that captured the main cellular processes. In cases of background noise, the Despeckle function was applied to improve image clarity. The resulting binary images were converted into skeletons using the Skeletonize plugin. Branching parameters were quantified using the Analyze Skeleton (2D/3D) tool.

For quantification of fluorescence intensity, Fluorescence images were processed and quantified using Fiji/ImageJ software (NIH, USA). The mean fluorescence intensity for each channel was measured within the regions of interest (ROIs) using the Measure function in Fiji. A consistent automatic thresholding method was applied to segment the signal from the background. Regions outside the signal, including scale bars and artifacts, were excluded. Mean fluorescence intensity was measured within the threshold-defined area. All images were acquired under identical microscope settings to ensure comparability.

### 2.9 Western Blotting

Protein Assay Dye Reagent Concentrate (Bio-Rad, USA) was used to measure the concentration of extracted proteins following the manufacturer’s instructions. For each sample, 15ug protein was mixed with 4× SDS loading buffer (Bio-Rad, USA), heated at 95□ for 5□minutes, and separated by 4–20% Mini Protean TGX gels (Bio-Rad, USA). Sample transfer to 0.2□μm nitrocellulose membranes (Amersham™ Protran™, Cytiva, Germany) using the Fisherbrand™ Semi-Dry Blotter (Thermo Fisher Scientific, UK) was performed according to the manufacturer’s protocol. Membranes were then blocked with 5% nonfat milk (Cell signaling technology, USA) Tris-buffered saline with Tween-20 (TBST) for 1□hour at room temperature, followed by overnight incubation at 4□°C with primary antibodies. After three washes with TBST, membranes were incubated with species-appropriate horseradish peroxidase (HRP)-conjugated secondary antibodies for 1□hour at room temperature. The details of primary and secondary antibodies are listed in Table S2.

Signal detection was performed using Clarity™ Western ECL substrate (Bio-Rad, USA), and membranes were imaged with ChemiDoc^TM^ MP imaging system (Bio-Rad, USA). Band intensities were quantified using ImageJ software (version 1.54) and normalized to loading control proteins.

### 2.10 Enzyme-linked immunosorbent assays (ELISA)

Mouse serum samples and mouse microglia cell culture supernatants were used to measure HMOX2 with a commercial HMOX2 kit (MyBiosourse, USA) according to manufacturer’s instructions. Mouse serum samples were also used to measure IL-6 and IL-18 level using IL-6 and IL-18 mouse Elisa kits (R&D systems, USA). Optical density (OD) values were measured using the SpectraMax microplate reader (Molecular Devices, USA).

### 2.11 Cell culture and measurement of ROS production via 2**′**,7**′**-dichlorodihydrofluorescein diacetate (DCFH-DA) assay

The mouse microglia cell line SimA9 cells were cultured in DMEM/F12 GlutaMAX™ culture media (Gibco, UK), supplemented with 10% fetal bovine serum (FBS, Sigma-Aldrich, USA), 5% horse serum (Sigma-Aldrich, USA), and 1% penicillin–streptomycin (Gibco, UK). Cells were maintained either in 6-well plates (1.5×10□/well) or in 8-chamber polystyrene tissue culture–treated glass slides (1× 10^4^/chamber). Prior to treatment with H□O□, IL-6, or IL-18, cells were preconditioned in Opti-MEM (Gibco, UK) overnight. They were then treated with 100 μM hydrogen peroxide (H□O□, Merck, Germany), 20 ng/mL IL-6 (R&D systems, USA), or 50 ng/mL IL-18 (MedChemExpress, USA) in Opti-MEM for 4, 8, or 24 h.

After treatment, culture supernatants from the 6-well plates were collected for subsequent HMOX2 ELISA measurements. And proteins from cells were extracted for western blot. For ROS detection, cells seeded in 8-chamber slides were washed with PBS and incubated with 10 μM DCFH-DA (Sigma-Aldrich, USA) for 30 min at 37 °C. The probe solution was then removed, cells were washed with PBS, and fluorescence signal was acquired using a Zeiss LSM 880 confocal microscope. Fluorescence intensity was quantified using ImageJ software (version 1.54).

### 2.12 Statistical analysis

For the analyses of human plasma proteomics data, multiple paired t-test was performed to assess differences in expression between sJIA samples (n=16) and healthy controls (n =□16). P values were then adjusted for multiple testing using the Benjamini–Hochberg false discovery rate (FDR) method. To compare categorical variables such as sex ratio, and treatment distribution between the two sJIA clusters, defined based on the neurobiological protein profile, Fisher’s exact test was used. Mann–Whitney U test was used to determine the differences in age and disease duration between two sJIA clusters. Spearman correlation analyses with the Benjamini–Hochberg adjustment were used to determine correlations between target proteins and selected clinical parameters. Multiple paired t-tests with Holm–Bonferroni adjustment were used to assess changes between the active and inactive phases in paired samples from sJIA patients (n = 12) and HCs (n = 12). Experimental data from mice were analyzed with an unpaired t-test. Spearman correlation analysis was performed to identify the correlations between HMOX2/IL-6/IL-18 levels in mouse serum samples and CNS changes. RNA sequence data was analyzed according to the protocol in RNA sequencing and data analysis section. Statistical analysis was performed using GraphPad Prism 9.0.0 (GraphPad Software, USA) or R software (v4.4.1).

For statistical analyses of the Olink profiling data, an FDR-adjusted p-value < 0.05 was considered statistically significant. For the RNA-seq analysis, genes with FDR-adjusted p-value < 0.1 and |log2 Fold Change| > 0.585 were considered significant. For analyses involving a single predefined comparison without multiple testing in sJIA patients and mice, a p-value < 0.05 was considered statistically significant. For multiple comparisons within the sJIA cohort, an adjusted p-value < 0.05 was considered statistically significant.

## 3. Results

### 3.1 sJIA patients exhibit an altered neurobiological protein profile in plasma compared with healthy individuals

We collected plasma samples from patients with active sJIA and age- and sex-matched HCs, and profiled neurobiological related proteins using the Olink Neuro Exploratory panel (Figure 1A). Three-dimensional PCA revealed a clear separation between HCs and sJIA patient (Figure 1B). Hierarchical cluster analysis also demonstrated a distinct separation between the two groups and the relative abundance of each protein across individuals was visualized in a heatmap (Figure 1C). Overall, 32 proteins were significantly downregulated in sJIA patients compared to HCs, while one protein, progestagen associated endometrial protein (PAEP) was upregulated (Figure 1D, Table S3).

**Figure 1.**
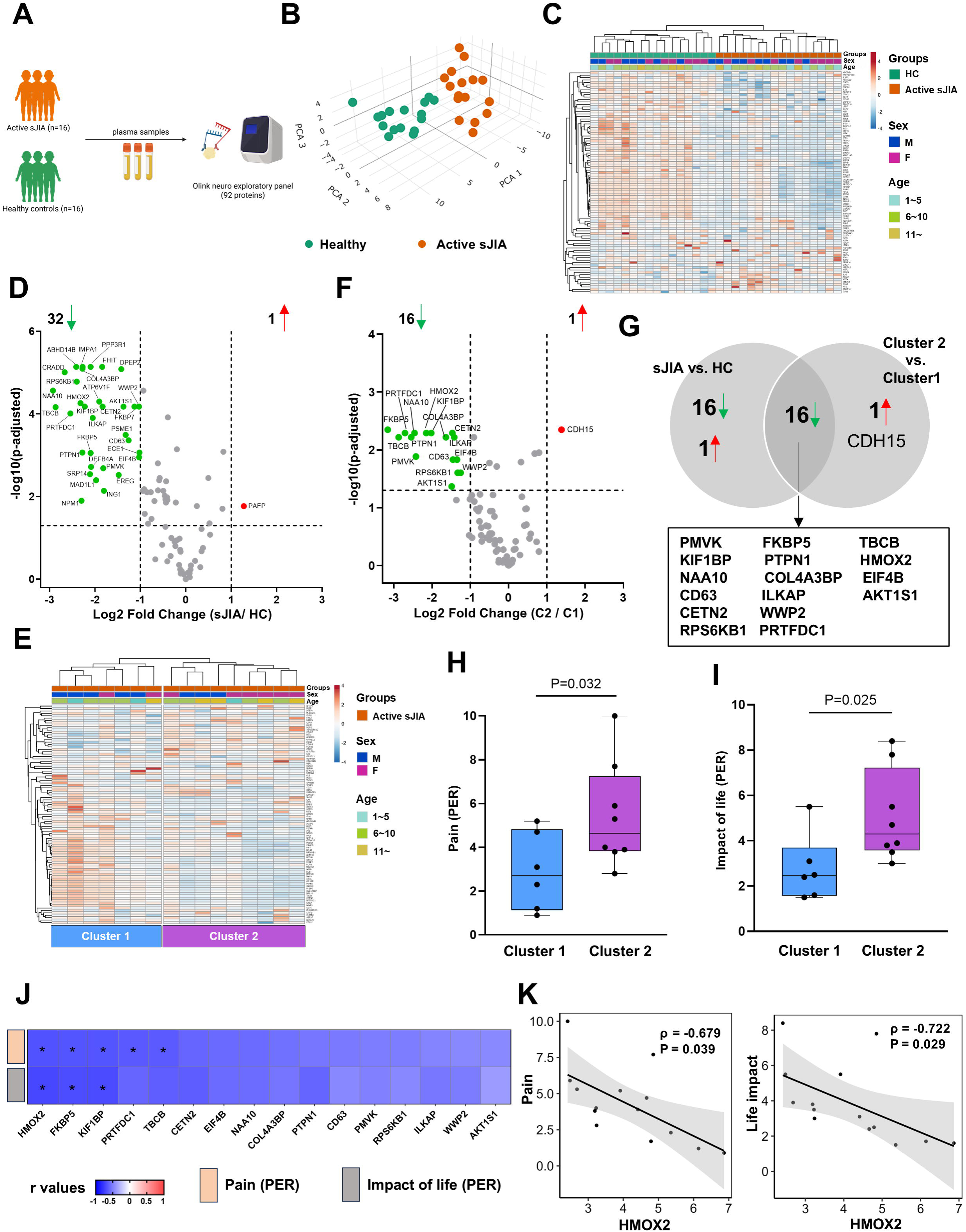
sJIA patients exhibit an altered neurobiological protein profile, which is associated with higher pain and greater life impact**. (A)** Schematic diagram of a cross-sectional study profiling neurobiological related proteins, including 16 active sJIA, and 16 age- and sex-matched HCs. **(B)** PCA plot of neuro-related plasma proteins showing separation between sJIA patients and HCs. Each point represents one subject (n = 32; 16 active sJIA, and 16 HCs). **(C)** Heatmap of protein expression in sJIA and HCs, with hierarchical clustering revealing separation between sJIA and HCs. **(D)** Volcano plot of DEPs in sJIA vs. HCs. Dashed lines indicate significance thresholds. **(E)** Unsupervised hierarchical clustering analysis grouping sJIA patients into two clusters based on their neurobiological protein expression patterns. **(F)** Volcano plot of DEPs in Cluster 2 vs. Cluster 1. Dashed lines indicate significance thresholds. **(G)** Venn diagram showing overlapping DEPs between sJIA vs. HCs and Cluster 2 vs. Cluster 1. **(H)** Box plot comparing pain scores between active sJIA patients in Cluster 1 and Cluster 2. One patient in each cluster is missing pain score data. **(I)** Box plot comparing life impact scores between active sJIA patients in Cluster 1 and Cluster 2. One patient in each cluster is missing life impact score data. **(J)** Heatmap of correlations between DEPs and pain/life impact in active sJIA patients. **(K)** Correlation analysis showing HMOX2 as the strongest negatively correlated protein with pain and life impact in active sJIA patients, illustrated by scatter plot. Statistics: (D) paired t-test with Benjamini–Hochberg correction for multiple comparisons; (F) unpaired t-test with Benjamini–Hochberg correction for multiple comparisons; (H, I) unpaired t-test; (J, K) Spearman correlation analysis with Benjamini–Hochberg correction. *: *P* < 0.05.

### 3.2 The altered neurobiological protein profile links to higher pain and life impact in sJIA patients and associated with disease duration

To further explore neurobiological proteomic alterations in sJIA, we performed unsupervised hierarchical clustering based on proteomic profiles from sJIA patients. This analysis revealed two distinct clusters in sJIA, comprising 7 patients in Cluster 1 and 9 patients in Cluster 2. (Figure 1E). Interestingly, the differences in protein expression between these two clusters mirrored those observed when comparing sJIA with HCs. Specifically, in cluster 2, a total of 16 proteins were downregulated compared with cluster 1, and these proteins also showed decreased level in the overall comparison between sJIA and HCs. One protein, cadherin-15 (CDH15) was upregulated in cluster 2 as compared with cluster 1 (Figure 1F–1G, Table S4).

To assess the clinical significance of these proteomic changes, we compared clinical parameters reflecting neurobiological well-being between the two clusters. As psychometric data was unavailable for the sJIA cohort, patient-reported pain and life impact scores were used as surrogate indicators of neurological status. Patients in Cluster 2 reported higher pain and life impact scores compared to those in Cluster 1 (Figure 1H–1I). We then examined the associations between 16 downregulated proteins (from Figure 1G) and patient-reported pain and life impact scores in sJIA. Correlation analysis identified 11 proteins negatively associated with pain and 5 proteins negatively associated with life impact in active sJIA patients (Figure 1J). Among these, HMOX2 showed the strongest negative correlation (lowest adjusted-p value) with both pain (Spearman r=-0.679, *P_adj_*=0.039) and life impact (Spearman r=-0.722, *P_adj_*=0.029) (Figure 1K, Table S5-S6). These results indicate that the altered neurobiological profile may reflect underlying neurofunctional impairments and suggest mechanisms potentially affecting psychiatric well-being.

To determine the factors influencing clustering based on neurobiological profiles, we compared the demographics and disease characteristics of sJIA patients in the two clusters, including sex, age at onset, age at sampling, disease duration, disease activity, C-reactive protein (CRP) measurements, autoantibodies, and treatment. Notably, patients in Cluster 2 had a substantially longer disease duration (Table S7). No other significant differences were observed between the two clusters, and treatment regimens did not differ, suggesting that these factors were unlikely to account for the observed clustering (Figure S1A, Table S7).

### 3.3 Altered neurobiological protein profiles persist during the inactive phase of sJIA

Among the 16 active sJIA patients, 12 also had plasma samples collected later during the inactive phase. We compared neurobiological profiles between paired samples from these patients, with age- and sex-matched HCs serving as a baseline (Figure 2A). PCA revealed that inactive sJIA samples were, unexpectedly, more distinct from HCs than active sJIA samples (Figure 2B). Interestingly, all proteins previously identified as negatively correlated with pain and life impact were even lower in the inactive phase compared with the active phase (Figure 2C). These findings suggest that even after clinical remission of active sJIA, neurobiological alterations may continue to progress.

**Figure 2.**
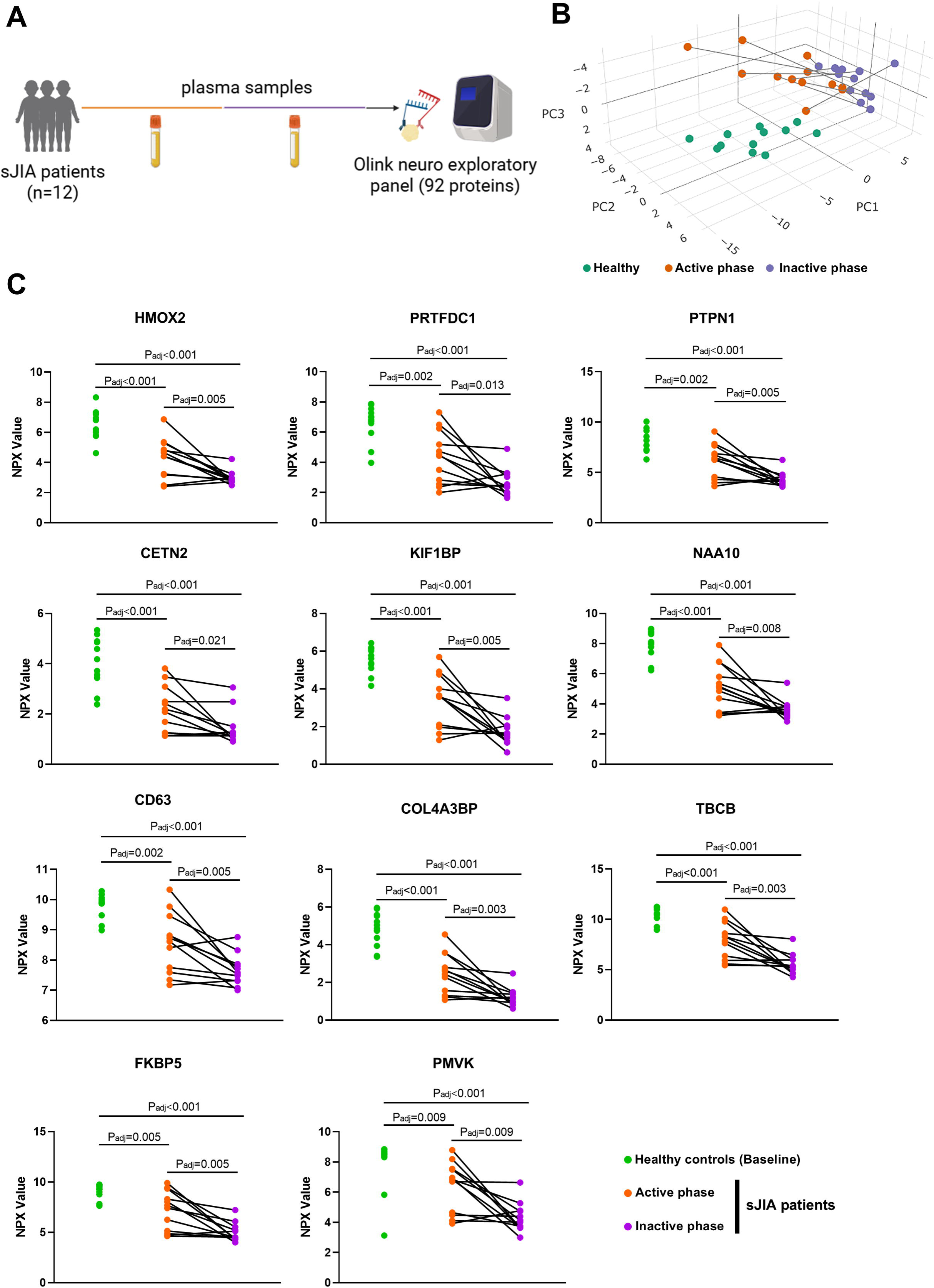
Paired comparison of neurobiological protein profile changes in 12 sJIA patients across active and inactive phases, with 12 age- and sex-matched HCs. **(A)** Schematic diagram of a paired study profiling neurobiological related proteins, including 12 sJIA patients with samples from both active and inactive phases, and 12 age- and sex-matched HCs. **(B)** PCA plot of neurobiological related plasma proteins showing the distribution of sJIA patients (active vs inactive) and HC. Each point represents one subject, with lines connecting paired active and inactive samples from the same patient (n = 36; 12 active sJIA, 12 inactive sJIA, and 12 HCs). **(C)** Line plot showing longitudinal changes in proteins identified as related to clinical parameters, from active to inactive sJIA, using age- and sex-matched healthy individuals as the baseline. Statistics: (C) Multiple paired t-tests with Holm–Bonferroni adjustment.

### 3.4 Neuroinflammation and impaired neurogenesis in mice with chronic arthritis

As no animal model currently fully recapitulates the complex features of sJIA, we employed the CIA model and maintained mice for 3 months to mimic features of chronic autoimmune arthritis and to further investigate the link between arthritis and neurobiological alterations (Figure 3A). The arthritis score peaked on day 69 and subsequently fluctuated, resembling the flare and remission pattern observed in patients (Figure 3B). Arthritis parameters were quantified for each mouse (Figure S2A).

**Figure 3.**
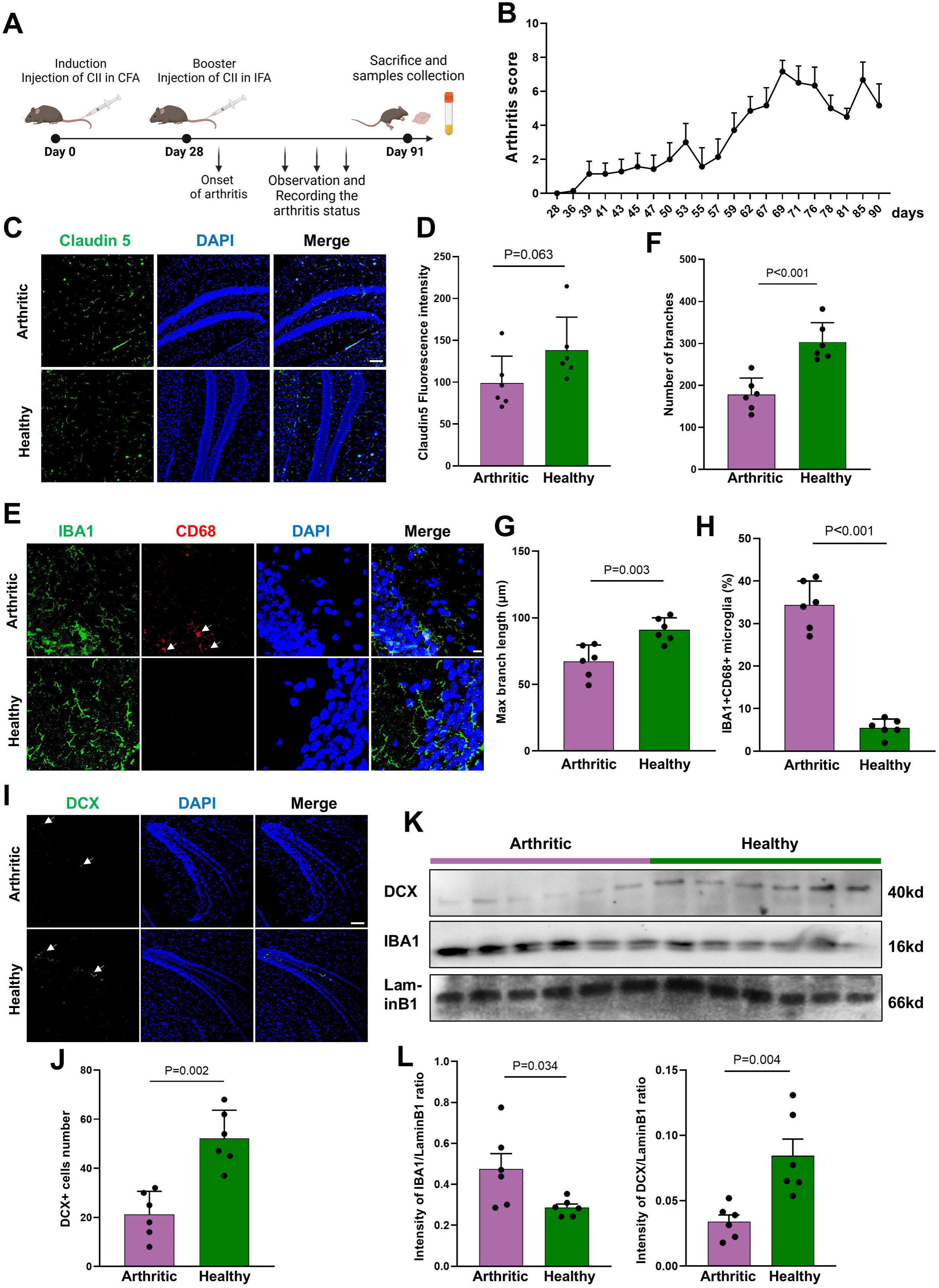
Chronic arthritic mice exhibit neuroinflammation and impaired neurogenesis in the hippocampus. **(A)** Schematic diagram of the chronic arthritis mouse model (collagen-induced arthritis, CIA). **(B)** Line plot showing changes in arthritis scores of immunized mice over the disease course. **(C)** Representative images of immunofluorescence (IF) staining for Claudin-5 in the hippocampus of arthritic (n=6) and healthy (n=6) mice. Scale Bar: 100μm. **(D)** Box plots quantifying Claudin-5 fluorescence intensity by IF in arthritic and healthy mice. **(E)** Representative images of IF staining for microglial markers (IBA1 and CD68) in the hippocampus of arthritic (n=6) and healthy (n=6) mice. Scale Bar: 10μm. **(F–G)** Box plots quantifying microglial morphology (branch number and branch length) in arthritic and healthy mice. **(H)** Box plots showing the number of activated microglial cells (IBA1D CD68D) by IF in arthritic and healthy mice. **(I)** Representative images of IF staining for DCX in the hippocampus of arthritic (n=6) and healthy (n=6) mice. Scale Bar: 100μm. **(J)** Box plots quantifying the number of DCX+ neurons by IF in arthritic and healthy mice. **(K–L)** Western blotting results showing downregulation of DCX and upregulation of IBA1 in arthritic hippocampus compared with healthy controls. Statistics: (D, F, G, H, J, L) unpaired t-test.

We focused our analysis on the hippocampus and cortex, regions known to be critical for cognition and mood regulation [26]. In the hippocampus, we investigated Claudin-5 expression, a tight junction protein critical for maintaining BBB integrity, but did not observe significant changes between arthritic and healthy mice, suggesting no apparent BBB impairment (Figure 3C-3D). In contrast, microglia in arthritic mice exhibited a pronounced amoeboid morphology, characterized by fewer and shorter branches (Figures 3E–3G). Activated microglia (IBA1□CD68□) were significantly increased in the hippocampus of arthritic mice, indicating neuroinflammation (Figures 3E, 3H). Furthermore, neurogenesis appeared impaired, as evidenced by a reduction in DCX□ neuronal cells in the hippocampus of arthritic mice (Figure 3I-3J). These observations were further validated by western blot analysis (Figures 3K–3L).

In contrast, no significant changes were observed in the cortex between arthritic and healthy mice. Claudin-5 expression in the cortex showed no notable difference (Figures S2B–S2C). Similarly, microglial activation, assessed by IBA1 and CD68 staining, did not show significant alterations in the cortex (Figures S2D–S2F).

### 3.5 Mitochondrial dysfunction and oxidative stress are important contributors to hippocampal alterations in experimental chronic arthritis

To further elucidate the molecular mechanisms underlying neuroinflammation, BBB integrity and decreased neurogenesis, as suggested by immunofluorescence analysis of the hippocampus of arthritic mice, we isolated hippocampal tissues from both arthritic and healthy control mice for transcriptomic profiling using RNA sequencing, with sequence quality assessed by FastQC and MultiQC analysis, including duplicate read analysis, per-base and per-sequence quality metrics, and per-sequence GC content (Fig S3A-S3D).

PCA plot of the hippocampal transcriptomes revealed a separation between arthritic and control groups (Figure 4A), indicating substantial transcriptional changes in the hippocampus during arthritis. Differential gene expression analysis identified 94 upregulated and 517 downregulated genes in the hippocampus of arthritic mice as compared with controls (Figure 4B). We performed gene set enrichment analysis to explore dysregulated pathways. Pathways related to neurogenesis (e.g., Positive regulation of neural precursor cell proliferation, and forebrain neuron development), BBB integrity (e.g., Establishment of endothelial barrier, and extracellular structure organization) were significantly downregulated in arthritic mice (Figure S4A), supporting the observed hippocampal dysfunction and aligning with our histological findings.

**Figure 4.**
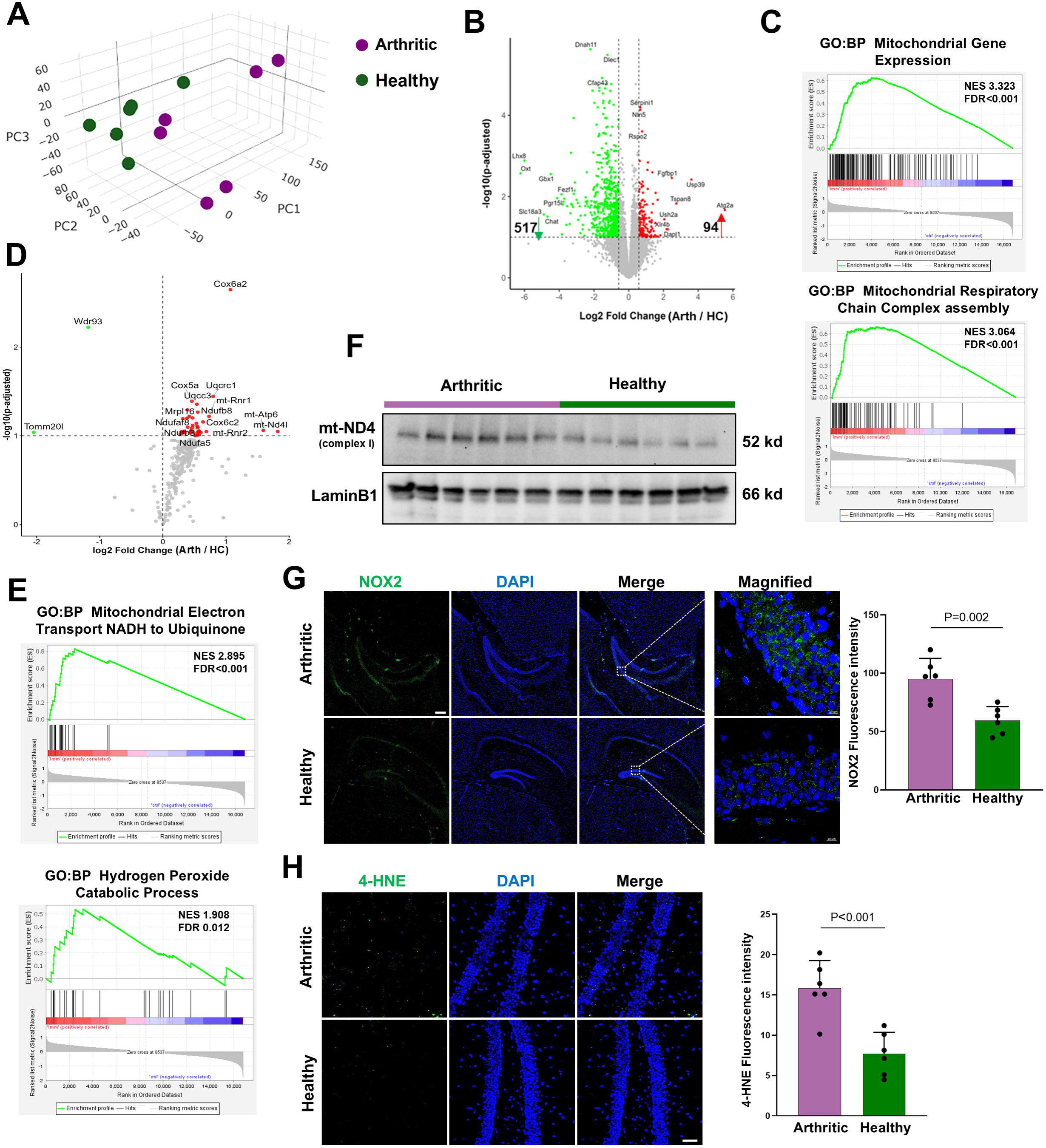
RNA-seq analysis identifies mitochondrial dysfunction and oxidative stress associated with hippocampal neuroinflammation in arthritic mice. **(A)** PCA plot of RNA-seq data showing separation between arthritic (n = 6) and healthy (n = 6) mice. **(B)** Volcano plot displaying 94 upregulated and 517 downregulated genes in the hippocampus of arthritic mice compared with healthy mice. Dashed lines indicate significance thresholds. **(C)** GSEA plots showing activation of pathways related to mitochondria and the mitochondrial respiratory chain in arthritic mice. **(D)** Volcano plot showing mitochondrial-related DEGs in the hippocampus of arthritic mice compared with healthy controls. Dashed lines indicate significance thresholds. **(E)** GSEA plots identifying activation of pathways involving mitochondrial complex I and the hydrogen peroxide catabolic process in arthritic mice. **(F)** Western blot results showing upregulation of mt-ND4 in arthritic hippocampus compared with healthy controls. **(G)** Representative images of IF staining for NOX2 in the hippocampus of arthritic (n = 6) and healthy (n = 6) mice, with corresponding box plot quantifying NOX2 fluorescence intensity. Scale Bar: 200μm. **(H)** Representative images of IF staining for 4-HNE in the hippocampus of arthritic (n = 6) and healthy (n = 6) mice, with corresponding box plot quantifying 4-HNE fluorescence intensity. Scale Bar: 50μm. Statistics: (B, D) RNA-seq counts normalized and differential expression calculated using the DESeq2 package; (G, H) unpaired t-test.

Interestingly, GSEA revealed that the predominant upregulated pathways in the hippocampus of arthritic mice were related to mitochondrial function and the mitochondrial respiratory chain (Figure 4C, Table S8). Based on this observation, we hypothesized that mitochondrial dysfunction may contribute to neuroinflammation in the hippocampus. To explore this possibility, we curated a list of mitochondria-related genes (based on GO:0005739 dataset) and identified differentially expressed genes (DEGs, P_adj_ < 0.1). In total, 40 mitochondria-associated genes were upregulated, while only 2 were downregulated in the hippocampus of arthritic mice (Figure 4D). Notably, many of these DEGs were components of mitochondrial complex I (e.g., mt-ND4L, Ndufaf8, Ndufb3, Ndufa5, and Ndufb8). Mitochondrial complex I is an important site of ROS production and has previously been reported to exhibit increased activity in microglia, thereby sustaining neuroinflammation in chronic inflammatory disorders[27, 28]. Our GSEA results also demonstrated that pathways related to mitochondrial complex I and oxidative stress were significantly enriched in the hippocampus of arthritic mice (Figure 4E). Therefore, we investigated mitochondrial complex I activity and oxidative stress in the hippocampus of arthritic and healthy mice. Immunofluorescence staining showed abundant complex I expression in both groups, with no significant differences detected (Figure S4B), possibly due to high baseline expression in hippocampal tissue. However, western blot analysis revealed increased mt-ND4 protein levels in the hippocampus of arthritic mice (Figure 4F, Figure S4C). Moreover, NADPH oxidase 2 (NOX2), a key contributor to ROS production, was markedly upregulated in the hippocampus of arthritic mice (Figure 4G). Additionally, immunostaining for 4-hydroxynonenal (4-HNE), a lipid peroxidation marker of oxidative stress, also revealed increased levels in the arthritic group (Figure 4H). These findings indicate elevated oxidative stress in the hippocampus of arthritic mice and involved in neuroinflammation during chronic arthritis.

### 3.6 HMOX2 is downregulated in arthritic mice, and serum HMOX2 levels are negatively correlated with hippocampal neuroinflammation

In the analyses of sJIA, HMOX2 emerged as a protein of particular interest. HMOX2 has a well-established role in oxidative stress regulation and our RNA-seq data emphasized the involvement of oxidative stress in the hippocampus of arthritic mice. More importantly, its plasma expression was markedly downregulated in patients with sJIA and demonstrated the strongest negative correlation (Figure 1J-1K) with clinical measures of pain and life impact in active sJIA patients. Given its tissue-enriched expression in the brain and testis[29], these findings prompted us to further investigate a potential biomarker role of HMOX2 for neuroinflammation associated with chronic arthritis.

We first examined HMOX2 expression in hippocampal tissue. Immunofluorescence staining revealed that HMOX2 was abundantly expressed in hippocampus of healthy mice, with prominent localization in homeostatic microglia, as demonstrated by co-localization with P2Y purinergic receptor 12 (P2ry12), a marker of homeostatic microglia (Figure 5A-5B). In contrast, HMOX2 expression was significantly decreased in the hippocampus of arthritic mice (Figure 5C–5E). Consistently, serum levels of HMOX2 were also markedly reduced in arthritic mice (Figure 5F). Importantly, serum HMOX2 levels were negatively correlated with microglial activation in the hippocampus, as indicated by the frequency of IBA1□CD68□ microglia (Figure 5G). Given that HMOX2 also exhibited a strong negative correlation with pain severity and life impact scores in sJIA patients (Figure 1J-1K), peripheral HMOX2 may serve as a promising biomarker reflecting neurobiological alterations in the brain during chronic arthritis.

**Figure 5.**
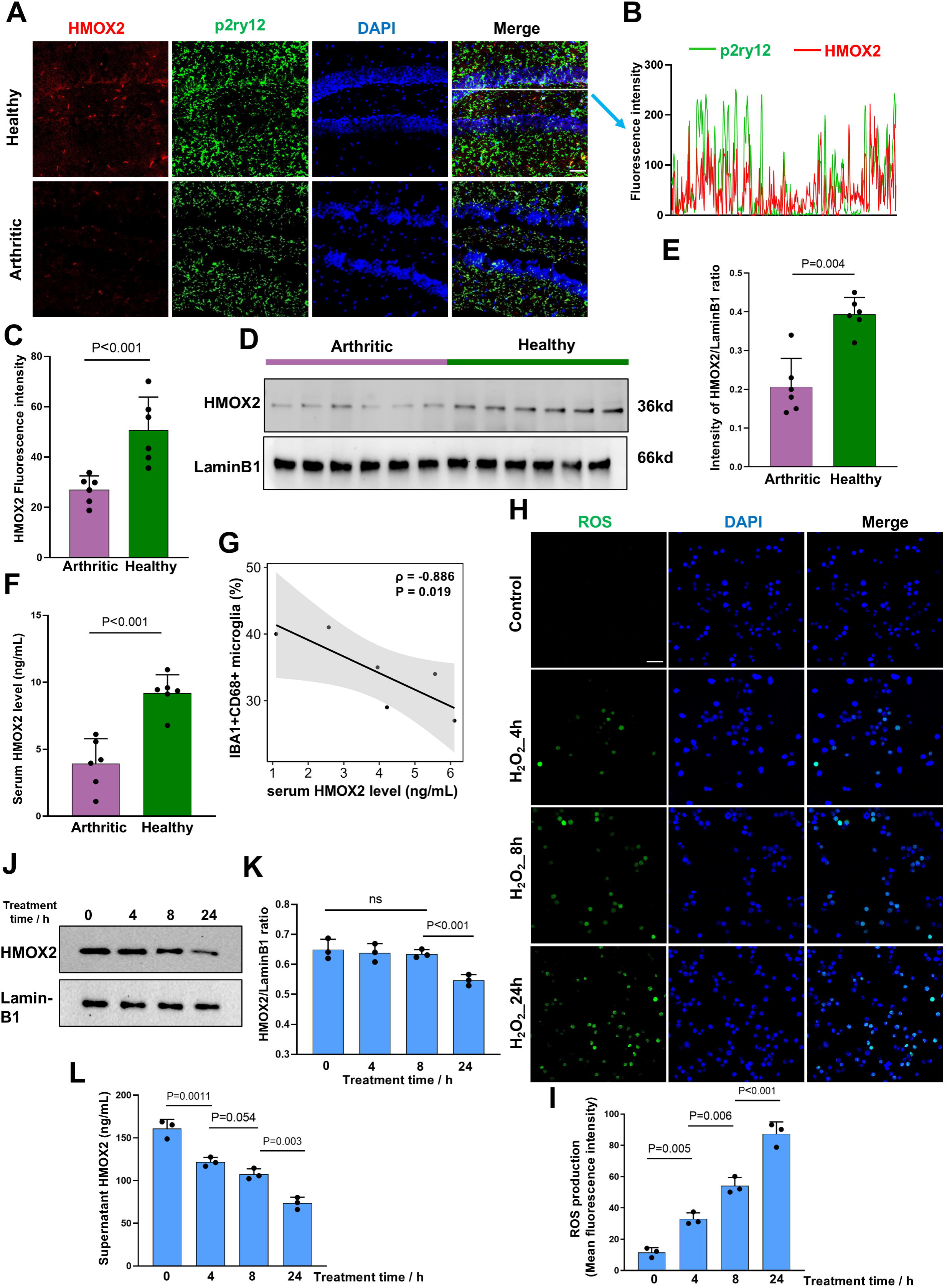
HMOX2 is decreased in the hippocampus and serum of arthritic mice, and its downregulation is associated with microglial oxidative stress. **(A)** Representative images of IF staining for HMOX2 and the homeostatic microglial marker P2ry12 in the hippocampus of arthritic (n = 6) and healthy (n = 6) mice. Scale Bar: 50μm. **(B)** Intensity profiles of HMOX2 (red) and P2ry12 (green) along the line indicated in Figure 4A in hippocampal tissue. **(C)** Box plots quantifying HMOX2 fluorescence intensity by IF in arthritic and healthy mice. **(D–E)** Western blot results showing decreased HMOX2 in the hippocampus of arthritic mice compared with healthy controls. **(F)** ELISA results showing reduced serum HMOX2 levels in arthritic mice compared with healthy controls. **(G)** Correlation analysis showing a negative correlation between serum HMOX2 and microglial activation in arthritic mice. **(H–I)** Representative images of DCFH-DA staining showing ROS levels in SimA9 cells treated with HDOD for 4, 8, and 24 h, with corresponding quantification of fluorescence intensity. Scale bar: 50μm. **(J–K)** Western blot results showing intracellular HMOX2 changes in SimA9 cells treated with HDOD for 4, 8, and 24 h, with corresponding quantification of grayscale intensity. **(L)** ELISA results showing downregulation of extracellular HMOX2 in SimA9 cells following HDOD treatment. Statistics: (C, E, F) unpaired t-test; (G) Spearman correlation analysis; (I, K, L) three independent experiments, one-way ANOVA followed by Tukey’s multiple comparisons test to assess differences among groups.

Based on the evidence of oxidative stress in the hippocampus of arthritic mice and reduced HMOX2 expression observed in both murine and clinical samples, we sought to explore the mechanistic link between peripheral HMOX2 and neuroinflammation. To this end, we conducted *in vitro* experiments using SimA9, a spontaneously immortalized mouse microglial cell line[30] selected due to our findings that HMOX2 is expressed in homeostatic microglia in mouse hippocampus. To mimic oxidative stress conditions in microglia, we treated SimA9 cells with H□O□. ROS production was significantly elevated as early as 4 hours post-treatment, confirming the induction of oxidative stress (Figure 5H-5I). Western blot analysis revealed that intracellular HMOX2 protein levels remained stable up to 8 hours but was decreased at 24 hours after H□O□ treatment (Figure 5J-5K). In contrast, ELISA results indicated a steady decline in extracellular HMOX2 levels during the whole period (Figure 5L). These findings suggest that oxidative stress reduces the release of HMOX2 from microglial cells. The reduction in circulating HMOX2 may be linked to systemic oxidative stress and indicate the development of hippocampal neuroinflammation.

### 3.7 IL-6 and IL-18 are key cytokines contributing to neuroinflammation during chronic arthritis

Previous studies have suggested that peripheral inflammatory cytokines may contribute to CNS neuroinflammation under pathological conditions such as autoimmune diseases, surgical trauma, and chronic infections [31–33]. To identify key cytokines potentially involved in neurobiological alterations during chronic arthritis, we profiled the inflammatory mediator landscape in the same sJIA samples using the Olink inflammation panel and performed a cross-panel correlation analysis between the two Olink datasets to evaluate the overlap and consistency of protein expression changes.

PCA plot demonstrated that active sJIA patients exhibited a distinct inflammatory signature, clearly separated from HCs (Figure 6A). Among the differentially expressed proteins, IL-6, IL-18, OSM, EN-RAGE, and MMP-1 were significantly elevated in active sJIA compared to healthy controls. Conversely, SCF, CD6, AXIN1, ST1A1, CXCL6, and CASP-8 were significantly decreased (Figure 6B, Table S9). We then compared the inflammation profiles between two sJIA clusters which were defined based on neurobiological protein profiles. IL-18 was significantly increased in cluster 2, while AXIN1 and SCF were decreased. We also noted a trend toward increased IL-6 levels in cluster 2 (P_adj_ = 0.054) (Figure 6C, Table S10). In addition, paired comparisons revealed that, unlike the continued progression of the neurobiological profile in the inactive phase, the inflammation profiles of sJIA patients partially returned to baseline when they were in inactive phase (Figure 6D).

**Figure 6.**
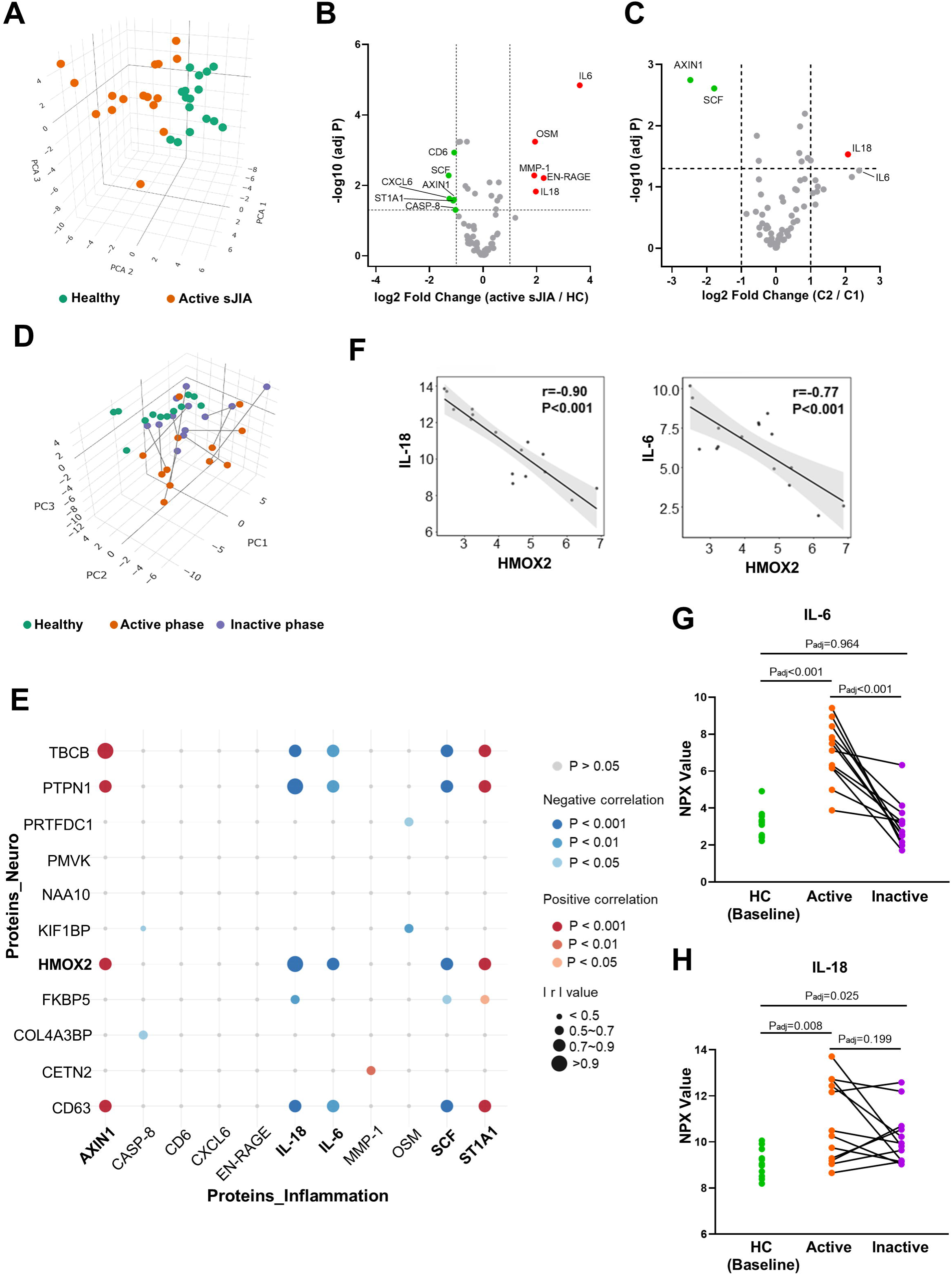
Inflammatory profiles of sJIA patients and neuro–inflammation co-analysis reveal correlations between HMOX2 and IL-6/IL-18. **(A)** PCA plot of inflammation-related plasma proteins showing separation between active sJIA patients and HCs. Each point represents one subject (n = 32; 16 active sJIA and 16 HCs). **(B)** Volcano plot of DEPs in active sJIA vs. HC. Dashed lines indicate significance thresholds. **(C)** Volcano plot of DEPs in Cluster 2 active sJIA vs. Cluster 1 active sJIA. Dashed lines indicate significance thresholds. **(D)** PCA plot of inflammation related plasma proteins showing the distribution of sJIA patients (active vs inactive) and HCs. Each point represents one subject, with lines connecting paired active and inactive samples from the same patient (n = 36; 12 active sJIA, 12 inactive sJIA, and 12 HCs). **(E)** Bubble plot showing correlations between DEPs from the inflammation panel and DEPs from the neuro panel in active sJIA patients. **(F)** Correlation analysis showing significant negative correlations between HMOX2 and IL-6/IL-18. **(G-H)** Line plot showing longitudinal changes of IL-6 and IL-18 from healthy to active to inactive sJIA. Statistics: (B) paired t-test with Benjamini–Hochberg correction for multiple comparisons; (C) unpaired t-test with Benjamini–Hochberg correction for multiple comparisons; (E, F) Pearson correlation analysis; (G, H) Multiple paired t-tests with Holm–Bonferroni adjustment.

To further explore the potential interplay between inflammatory proteins and neurobiological alterations, we performed correlation analyses between differentially expressed inflammatory proteins and neurobiological related DEPs associated with clinical measures of pain and life impact. In sJIA patients, IL-6, IL-18, and SCF exhibited significant negative correlations with HMOX2, whereas AXIN1 and ST1A1 were positively correlated with HMOX2 expression (Figure 6E-6F). We also investigated the IL-6 and IL-18 changes between active and inactive phase using the paired samples. We found IL-6 increased significantly in the active phase but returned to the baseline in inactive phase, while higher level of IL-18 could still be observed in inactive samples (Figure 6G-6H). Given the observed elevation of IL-6 and IL-18 in active sJIA patients and their negative association with HMOX2, we speculated IL-6 and IL-18 may play important roles in neuroinflammation during chronic arthritis.

To experimentally validate this hypothesis, we first assessed IL-6 and IL-18 levels in serum samples from mice with CIA. Consistent with our clinical observations, both IL-6 and IL-18 were significantly elevated in arthritic mice compared with healthy control mice (Figure 7A). Serum levels of IL-18 were negatively correlated with HMOX2 levels, while the correlation with IL-6 did not reach significance in arthritic mice (Figure 7B). But consistency with IL-18 and IL-6 patterns observed in sJIA patients were evident. We next examined the relationship between IL-6/IL-18 and microglial activation. Serum IL-6 levels did not show a positive correlation with microglial activation (Figure 7C). However, serum IL-18 levels exhibited a strong positive correlation with microglial activation (Figure 7C), suggesting a pronounced role for IL-18 in driving microglial inflammatory responses.

**Figure 7.**
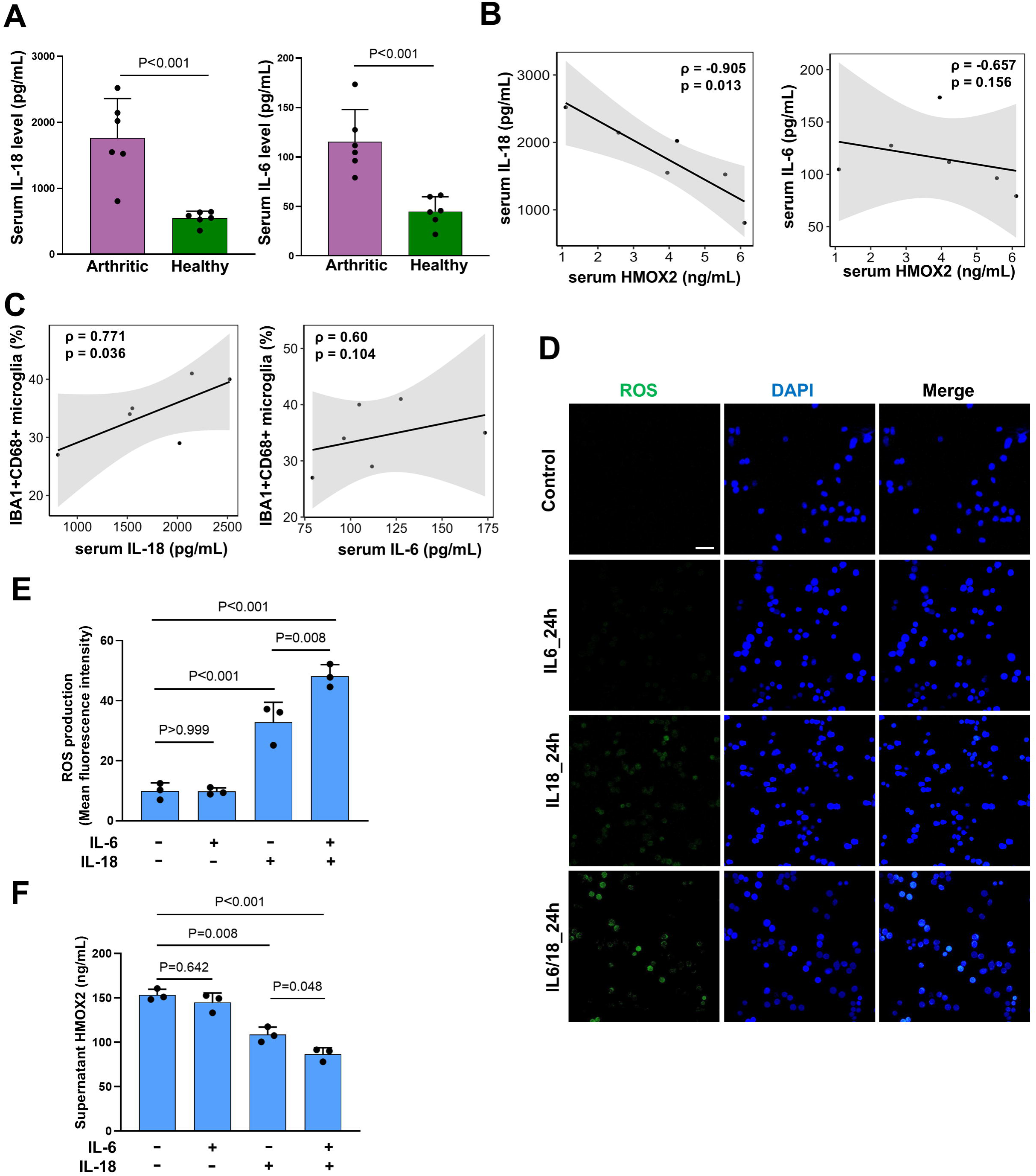
IL-6 and IL-18 are elevated in arthritic mice and synergistically trigger oxidative stress in microglia. **(A)** Box plots showing increased serum IL-6 and IL-18 levels in arthritic mice. **(B)** Correlation analysis showing a negative correlation between serum IL-18 and HMOX2, whereas IL-6 showed no significant correlation in arthritic mice. **(C)** Scatter plots showing the correlations between serum IL-18/IL-6 and microglial activation in arthritic mice. **(D–E)** Representative images of DCFH-DA staining showing ROS levels in SimA9 cells treated with IL-6, IL-18, or both for 24 h, with corresponding quantification of fluorescence intensity. Scale Bar: 50μm. **(F)** ELISA results showing downregulation of extracellular HMOX2 in SimA9 cells following IL-18 treatment or IL-6/IL-18 co-treatment. Statistics: (A) unpaired t-test; (B, C) Spearman correlation analysis; (E, F) three independent experiments, one-way ANOVA followed by Tukey’s multiple comparisons test to assess differences among groups.

To elucidate the direct effects of IL-6 and IL-18 on microglial oxidative stress, we treated SimA9 microglial cells with IL-6, IL-18, or a combination of both. IL-6 treatment alone did not induce significant ROS production, whereas IL-18 stimulation resulted in a moderate increase in ROS levels. Strikingly, co-treatment with IL-6 and IL-18 elicited a synergistic effect, leading to a substantial increase in ROS production (Figure 7D-7E). Corresponding ELISA assays for HMOX2 revealed no significant change in HMOX2 expression following IL-6 treatment, a moderate reduction with IL-18 treatment, and a more pronounced decrease upon combined IL-6/IL-18 stimulation (Figure 7F). Taken together, these findings suggest that IL-6 and IL-18, particularly IL-18, can synergistically induce oxidative stress in microglial cells, leading to downregulation of HMOX2 expression. This IL-6/IL-18-driven oxidative response may represent a potential mechanism underlying the neurobiological alterations observed during chronic arthritis (Figure 8).

**Figure 8.**
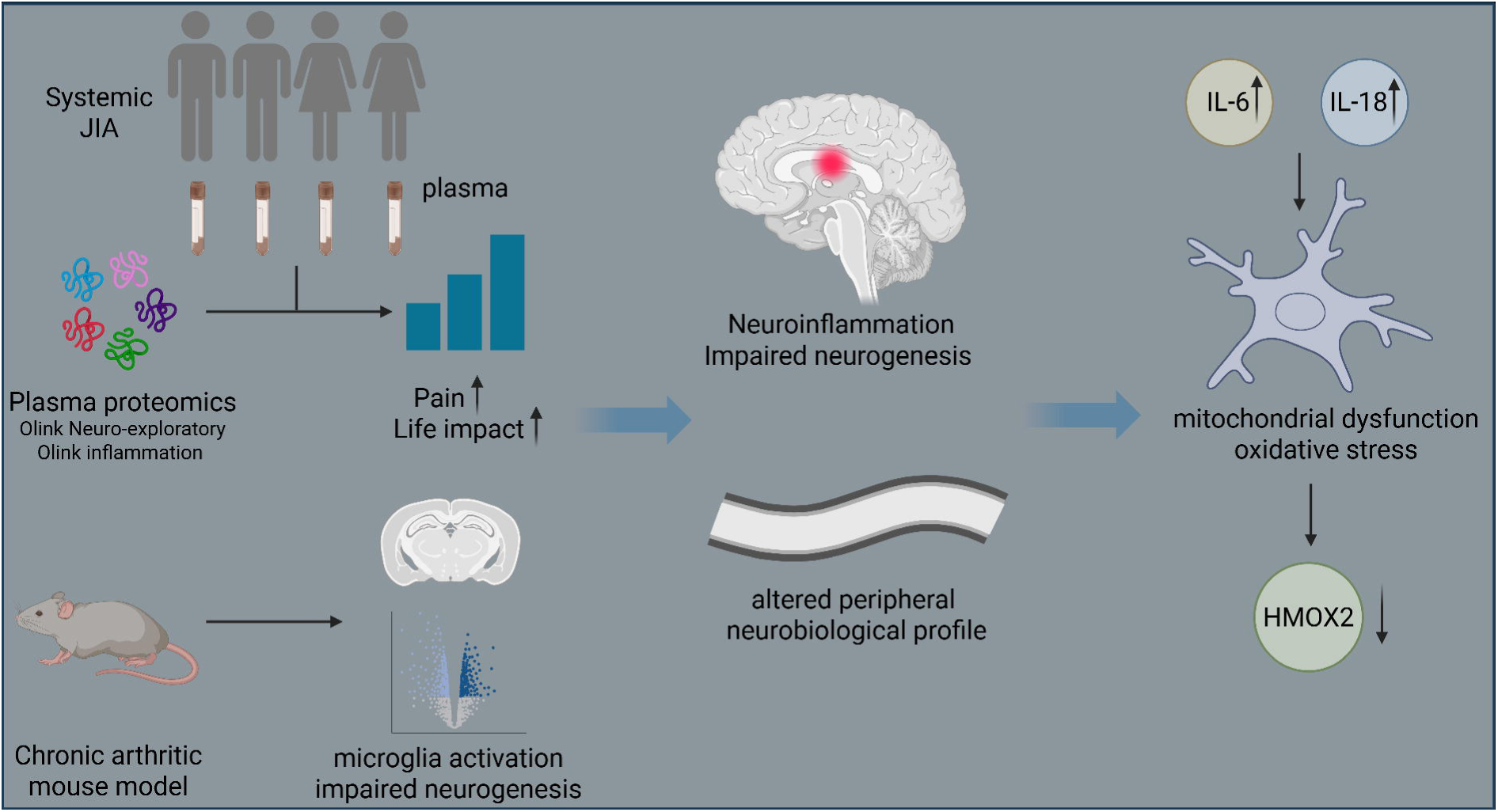
Graphic abstract of the mechanism underlying the neurobiological alterations during chronic arthritis.

## 4. Discussion

Over the decades, clinicians have increasingly recognized challenges related to emotional well-being and cognitive functioning in children living with JIA [34, 35]. However, current assessments of these neuropsychological features in JIA patients are not common [36], and direct evidence of structural or functional alterations within the CNS in JIA patients remains limited [37]. In this exploratory study, we demonstrate that patients with sJIA exhibit distinct neurobiological proteomic alterations in plasma. Reflecting to the clinical observations, we identified neuroinflammation and impaired neurogenesis in the hippocampus of arthritic mice, demonstrating CNS neuroinflammation is induced during chronic experimental arthritis. Lai et. al also reported change of the BBB permeability in CIA rats [38]. These findings provide molecular evidence supporting the recognized but mechanistically unexplained association between chronic arthritis and neuropsychiatric symptoms in individuals with JIA.

We identified an altered neurobiological protein profile in sJIA patients, with 32 proteins downregulated and 1 protein (PAEP) upregulated. Notably, a recent study on Prader–Willi syndrome found a similar pattern, with 24 out of 29 differentially expressed proteins decreased, including several overlapping proteins like HMOX2, NAA10, FKBP5, TBCB, and PTPN1[39]. These findings, along with a Mendelian randomization analysis by Reppeto et al., which linked lower plasma levels of approximately 75% of 184 proteins from the Olink Neurology and Olink Neuro Exploratory panels to increased risk of adverse outcomes with neuropsychiatric and non-neurological comorbidities [40], further suggest that reductions in circulating neurologically related proteins reflect biologically meaningful changes in the nervous system. We also identified two distinct neurobiological protein profile clusters among sJIA patients. Patients in Cluster 2 exhibited a more pronounced neurobiological profile, along with higher pain levels and greater life impact, compared to those in Cluster 1. This further supports that alteration in the neurobiological profile reflects the patients neurofunctional status to some extent. Further, our paired analysis revealed that even when patients transitioned from the active to the inactive phase, neurobiological alterations persisted, suggesting possible lasting effects on the nervous system. This could explain why patients in Cluster 2 have a longer disease duration. Consistently, a recent study reported sustained hippocampus neuroinflammation in JIA patients even during inactive disease[13].

Our study also highlights the critical role of elevated peripheral IL-6 and IL-18 in hippocampal neuroinflammation during chronic arthritis. IL-6 and IL-18 have been reported to contribute to neuroinflammation by promoting BBB disruption and astrocyte and microglial activation, and IL-18 also exerts potent effects on the CNS by amplifying neurotoxic cascades [41–43]. We observed positive correlations between serum IL-6 and IL-18 levels (the correlation with IL-6 did not reach statistical significance) and microglial activation in the hippocampus in arthritic mice, indicating that elevated levels of these cytokines could be contributing factors to the central neuroinflammatory processes detected. *In vitro* experiments further elucidated the distinct and synergistic effects of these cytokines. In sJIA, we found that both IL-6 and IL-18 levels were significantly elevated in patients with active disease. Notably, while IL-6 levels normalized following treatment, IL-18 levels remained persistently elevated in patients who had entered the inactive phase. Another study also reported elevated IL-18 levels in inactive sJIA patients, even among those receiving IL-1 inhibitors[44]. This residual elevation of IL-18 may partially explain the ongoing neurobiological protein profile changes observed in inactive sJIA patients, despite the apparent clinical remission of peripheral symptoms. Current therapeutic strategies primarily focus on IL-1 and IL-6 blockade in sJIA, which effectively controls systemic inflammation and joint manifestations[45]. However, the persistent elevation of IL-18 may represent a key driver of unresolved neurobiological alterations in these patients. Our study suggests that future therapeutic approaches should also consider targeting IL-18 to achieve a more comprehensive suppression of neuroinflammatory pathways during chronic arthritis.

HMOX2 was identified as a potential peripheral biomarker in our study, reflecting central neuroinflammation during chronic arthritis. HMOX2 is a heme oxygenase isoform that participates in heme degradation, redox regulation, and neuroprotection, and is enriched in the brain and testis[29, 46]. In this study, we consistently observed a significant reduction of peripheral HMOX2 levels in both sJIA patients and in arthritic mice. The expression of HMOX2 in hippocampus was also reduced in arthritic mice. Notably, plasma HMOX2 expression was inversely correlated with clinical assessments of pain intensity and life impact scores in patients. In arthritic mice, lower serum HMOX2 levels were associated with increased microglial activation in the hippocampus, indicating a relationship between peripheral HMOX2 expression and CNS neuroinflammatory status. We propose that a reduction in peripheral HMOX2 levels may reflect both systemic and CNS oxidative stress, thereby linking to neuroinflammation. Interestingly, in the APP/PS1 transgenic mouse model of Alzheimer’s disease, HMOX2 expression was also found to be decreased both in the hippocampus[47] and in serum[48], further supporting the relevance of HMOX2 in neuroinflammatory contexts. Based on these observations, peripheral HMOX2 levels may serve as a highly relevant indicator of CNS neuroinflammation in chronic arthritis.

Meanwhile, we observed that the pro-inflammatory cytokines IL-6 and IL-18 were elevated in arthritic mice and serum IL-18 levels positively correlated with microglial activation in the hippocampus. However, considering that IL-6 and IL-18 are pleiotropic cytokines and their upregulation is a common feature of various cell types and tissues [49, 50], their specificity for indicating CNS neuroinflammation is limited. In contrast, HMOX2 may confer a higher specificity for reflecting neuroinflammatory changes within the CNS. However, it should be noted that peripheral cells, such as endothelial cells, also express HMOX2[51]. Therefore, it is difficult to definitively attribute altered HMOX2 expression solely to microglial activation. Further investigation into the regulation of HMOX2 at both peripheral and central levels, as well as their potential association, is required.

Several limitations of this study should be acknowledged. First, the CIA mouse model, which is widely utilized to mimic Rheumatoid arthritis, was employed in this study. However, CIA is not an ideal model for sJIA. Given the current lack of an appropriate animal model that fully recapitulates sJIA pathology, we selected the CIA model to emphasize the chronicity of arthritis and its neuroinflammatory consequences, recognizing this as a necessary compromise. Additionally, age differences between the CIA model and the human sJIA population should also be considered as a potential limitation. Second, psychometric data were not available for the patient cohort, limiting our ability to directly evaluate neuropsychiatric outcomes. Consequently, we utilized pain and life impact scores as surrogate indicators of neurological involvement. However, it is important to note that these surrogate indicators are also influenced by a range of factors, including social and psychological factors[52], and may not exclusively reflect changes in the neurological system. Standardized psychiatric assessments and neuroimaging techniques such as functional MRI (fMRI), particularly targeting hippocampal activity are needed to enable a more objective and precise characterization of neurobiological alterations for future studies. Third, while our findings suggest a synergistic role of IL-6 and IL-18 in promoting neuroinflammation, these observations require further *in vivo* validation to confirm their mechanistic relevance and therapeutic potential.

In summary, this study identified alterations in neurobiological protein profiles in sJIA patients. And even in inactive phase, although inflammation and disease activity were reduced, persistent neurobiological changes remained, suggesting potential long-term implications for affected individuals. In chronic arthritic mice, we observed hippocampal neuroinflammation and increased oxidative stress. Notably, HMOX2 emerged as a potential peripheral biomarker of CNS neuroinflammation, with reduced levels significantly correlating with clinical measures of pain and life impact in patients, as well as with hippocampal microglial activation in mice. Furthermore, we found that IL-6 and, particularly, IL-18 play critical roles in neuroinflammation during arthritis, underscoring their relevance as a potential target for future therapeutic strategies.

## Contributors

Study conception and design: Xingzhao Wen, Heshuang Qu, Cecilia Aulin, and Helena Erlandsson Harris. Acquisition, analysis, and interpretation of the data: Xingzhao Wen, Malgorzata Benedyk-Machaczka, Daphne Chen, Cecilia Aulin, Erik Sundberg, Heshuang Qu, Maria Altman, and Helena Erlandsson Harris. Patient recruitment and sample collection: Erik Sundberg, Erik Melén, Cecilia Aulin, Maria Altman, and Helena Erlandsson Harris. Manuscript drafting and editing: Xingzhao Wen, Cecilia Aulin, and Helena Erlandsson Harris. Critical revision of the article for important intellectual content: Heshuang Qu, Malgorzata Benedyk-Machaczka, Daphne Chen, Erik Sundberg, Erik Melén, and Maria Altman. All authors read and approved the final version of the manuscript.

## Declaration of Interests

The authors declare no conflicts of interest.

## Acknowledgements

This work was supported by the Swedish Research Council (grant number 2021-02723), The Swedish Rheumatism Association, King Gustaf V 80-year Foundation and Region Stockholm (grant numbers 962319 and 9877921). We acknowledge the patients, as well as the data management and study teams involved in the JABBA Cohort. We thank Claudia Fredolini from SciLifeLab for her technical support with the proteomics analysis. We also thank the Genomics Core Facility (GCF) at the University of Bergen for their support with RNA sequencing.

## Data Sharing Statement

The raw RNA-sequencing data from mouse hippocampal tissues have been deposited in the Sequence Read Archive (SRA) under accession number PRJNA1348353. Other original data and materials are available from the corresponding author upon reasonable request.

## Notes

### Competing Interest Statement

The authors have declared no competing interest.

## References

1. Prince FH, Otten MH, van Suijlekom-Smit LW. Diagnosis and management of juvenile idiopathic arthritis. Bmj 2010; 341: c6434.

2. Martini A, Lovell DJ, Albani S, et al. Juvenile idiopathic arthritis. Nat Rev Dis Primers 2022; 8: 5.

3. Prakken B, Albani S, Martini A. Juvenile idiopathic arthritis. Lancet 2011; 377: 2138–2149.

4. Hinze CH, Foell D, Kessel C. Treatment of systemic juvenile idiopathic arthritis. Nat Rev Rheumatol 2023; 19: 778–789.

5. Kyllönen MS, Ebeling H, Kautiainen H et al. Psychiatric disorders in incident patients with juvenile idiopathic arthritis - a case-control cohort study. Pediatr Rheumatol Online J 2021; 19: 105.

6. Delcoigne B, Horne A, Reutfors J, Askling J. Risk of Psychiatric Disorders in Juvenile Idiopathic Arthritis: Population- and Sibling-Controlled Cohort and Cross-Sectional Analyses. ACR Open Rheumatol 2023; 5: 277–284.

7. Tezer D, Başay BK, Başay Ö et al. Cognitive performance, psychiatric comorbidities, and quality of life in pediatric patients with juvenile idiopathic arthritis: a comparative analysis with healthy controls. Child Neuropsychol 2024; 1–19.

8. Memari AH, Chamanara E, Ziaee V et al. Behavioral Problems in Juvenile Idiopathic Arthritis: A Controlled Study to Examine the Risk of Psychopathology in a Chronic Pediatric Disorder. Int J Chronic Dis 2016; 2016: 5726236.

9. Pedersen MJ, Høst C, Hansen SN et al. Psychiatric Morbidity Is Common Among Children With Juvenile Idiopathic Arthritis: A National Matched Cohort Study. J Rheumatol 2024; 51: 181–188.

10. Mena-Vázquez N, Ortiz-Márquez F, Cabezudo-García P et al. Longitudinal Study of Cognitive Functioning in Adults with Juvenile Idiopathic Arthritis. Biomedicines 2022; 10.

11. Mena-Vázquez N, Cabezudo-García P, Ortiz-Márquez F et al. Evaluation of cognitive function in adult patients with juvenile idiopathic arthritis. Int J Rheum Dis 2021; 24: 81–89.

12. Ding T, Hall A, Jacobs K, David J. Psychological functioning of children and adolescents with juvenile idiopathic arthritis is related to physical disability but not to disease status. Rheumatology (Oxford) 2008; 47: 660–664.

13. Han H, Weng Y, Yang S et al. Sustained hippocampal neuroinflammation and subsequent glutamatergic dysfunction in juvenile idiopathic arthritis: evidence from proton magnetic resonance spectroscopy ((1)H-MRS). Arthritis Res Ther 2025; 27: 232.

14. Han H, Weng Y, Liang H et al. Persistent neuroinflammation of the right insular cortex in children with juvenile idiopathic arthritis: a proton MRS study. Clin Rheumatol 2023; 42: 3059–3066.

15. Najjar S, Pearlman DM, Alper K et al. Neuroinflammation and psychiatric illness. J Neuroinflammation 2013; 10: 43.

16. Schroer AB, Ventura PB, Sucharov J et al. Platelet factors attenuate inflammation and rescue cognition in ageing. Nature 2023; 620: 1071–1079.

17. Felger JC, Lotrich FE. Inflammatory cytokines in depression: neurobiological mechanisms and therapeutic implications. Neuroscience 2013; 246: 199–229.

18. Haroon E, Miller AH, Sanacora G. Inflammation, Glutamate, and Glia: A Trio of Trouble in Mood Disorders. Neuropsychopharmacology 2017; 42: 193–215.

19. Nikolopoulos D, Manolakou T, Polissidis A et al. Microglia activation in the presence of intact blood-brain barrier and disruption of hippocampal neurogenesis via IL-6 and IL-18 mediate early diffuse neuropsychiatric lupus. Ann Rheum Dis 2023; 82: 646–657.

20. Alam A, Hana Z, Jin Z et al. Surgery, neuroinflammation and cognitive impairment. EBioMedicine 2018; 37: 547–556.

21. Zervides KA, Grenmyr E, Janelidze S et al. The impact of disease activity and interferon-α on the nervous system in systemic lupus erythematosus. Arthritis Res Ther 2025; 27: 60.

22. Qu H, Sundberg E, Aulin C et al. Immunoprofiling of active and inactive systemic juvenile idiopathic arthritis reveals distinct biomarkers: a single-center study. Pediatr Rheumatol Online J 2021; 19: 173.

23. Brand DD, Latham KA, Rosloniec EF. Collagen-induced arthritis. Nat Protoc 2007; 2: 1269–1275.

24. Ruth JH, Amin MA, Woods JM et al. Accelerated development of arthritis in mice lacking endothelial selectins. Arthritis Res Ther 2005; 7: R959–970.

25. Liberzon A, Birger C, Thorvaldsdóttir H et al. The Molecular Signatures Database (MSigDB) hallmark gene set collection. Cell Syst 2015; 1: 417–425.

26. Guise KG, Shapiro ML. Medial Prefrontal Cortex Reduces Memory Interference by Modifying Hippocampal Encoding. Neuron 2017; 94: 183–192.e188.

27. Okoye CN, Koren SA, Wojtovich AP. Mitochondrial complex I ROS production and redox signaling in hypoxia. Redox Biol 2023; 67: 102926.

28. Peruzzotti-Jametti L, Willis CM, Krzak G et al. Mitochondrial complex I activity in microglia sustains neuroinflammation. Nature 2024; 628: 195–203.

29. Intagliata S, Salerno L, Ciaffaglione V et al. Heme Oxygenase-2 (HO-2) as a therapeutic target: Activators and inhibitors. Eur J Med Chem 2019; 183: 111703.

30. Nagamoto-Combs K, Kulas J, Combs CK. A novel cell line from spontaneously immortalized murine microglia. J Neurosci Methods 2014; 233: 187–198.

31. Liu X, Nemeth DP, Tarr AJ et al. Euflammation attenuates peripheral inflammation-induced neuroinflammation and mitigates immune-to-brain signaling. Brain Behav Immun 2016; 54: 140–148.

32. Beltran-Velasco AI, Clemente-Suárez VJ. Impact of Peripheral Inflammation on Blood-Brain Barrier Dysfunction and Its Role in Neurodegenerative Diseases. Int J Mol Sci 2025; 26.

33. Becher B, Spath S, Goverman J. Cytokine networks in neuroinflammation. Nat Rev Immunol 2017; 17: 49–59.

34. Tezer D, Başay BK, Başay Ö et al. Cognitive performance, psychiatric comorbidities, and quality of life in pediatric patients with juvenile idiopathic arthritis: a comparative analysis with healthy controls. Child Neuropsychol 2025; 31: 692–710.

35. Fair DC, Rodriguez M, Knight AM, Rubinstein TB. Depression And Anxiety In Patients With Juvenile Idiopathic Arthritis: Current Insights And Impact On Quality Of Life, A Systematic Review. Open Access Rheumatol 2019; 11: 237–252.

36. Casaletto KB, Heaton RK. Neuropsychological Assessment: Past and Future. J Int Neuropsychol Soc 2017; 23: 778–790.

37. Upadhyay J, Lemme J, Cay M et al. A multidisciplinary assessment of pain in juvenile idiopathic arthritis. Semin Arthritis Rheum 2021; 51: 700–711.

38. Lai PH, Wang TH, Zhang NY et al. Changes of blood-brain-barrier function and transfer of amyloid beta in rats with collagen-induced arthritis. J Neuroinflammation 2021; 18: 35.

39. Pascut D, Giraudi PJ, Banfi C et al. Characterization of Circulating Protein Profiles in Individuals with Prader-Willi Syndrome and Individuals with Non-Syndromic Obesity. J Clin Med 2024; 13.

40. Repetto L, Chen J, Yang Z et al. The genetic landscape of neuro-related proteins in human plasma. Nat Hum Behav 2024; 8: 2222–2234.

41. Zhang W, Xiao D, Mao Q, Xia H. Role of neuroinflammation in neurodegeneration development. Signal Transduct Target Ther 2023; 8: 267.

42. Begum E, Mahmod MR, Rahman MM et al. IL-18 Blockage Reduces Neuroinflammation and Promotes Functional Recovery in a Mouse Model of Spinal Cord Injury. Biomolecules 2024; 15.

43. Kang R, Gamdzyk M, Lenahan C et al. The Dual Role of Microglia in Blood-Brain Barrier Dysfunction after Stroke. Curr Neuropharmacol 2020; 18: 1237–1249.

44. Quatrini L, Ciancaglini C, Caiello I et al. Innate Lymphoid Cell Phenotypic and Functional Alterations in Patients With Systemic Juvenile Idiopathic Arthritis. Arthritis Rheumatol 2025; 77: 1442–1450.

45. Fautrel B, Mitrovic S, De Matteis A et al. EULAR/PReS recommendations for the diagnosis and management of Still’s disease, comprising systemic juvenile idiopathic arthritis and adult-onset Still’s disease. Ann Rheum Dis 2024; 83: 1614–1627.

46. Muñoz-Sánchez J, Chánez-Cárdenas ME. A review on hemeoxygenase-2: focus on cellular protection and oxygen response. Oxid Med Cell Longev 2014; 2014: 604981.

47. Vidal C, Daescu K, Fitzgerald KE et al. Amyloid β perturbs elevated heme flux induced with neuronal development. Alzheimers Dement (N Y) 2019; 5: 27–37.

48. Xing S, Shen D, Chen C et al. Early induction of oxidative stress in a mouse model of Alzheimer’s disease with heme oxygenase activity. Mol Med Rep 2014; 10: 599–604.

49. Murakami M, Kamimura D, Hirano T. Pleiotropy and Specificity: Insights from the Interleukin 6 Family of Cytokines. Immunity 2019; 50: 812–831.

50. Ihim SA, Abubakar SD, Zian Z et al. Interleukin-18 cytokine in immunity, inflammation, and autoimmunity: Biological role in induction, regulation, and treatment. Front Immunol 2022; 13: 919973.

51. Chen RJ, Yuan HH, Zhang TY et al. Heme oxygenase-2 suppress TNF-α and IL6 expression via TLR4/MyD88-dependent signaling pathway in mouse cerebral vascular endothelial cells. Mol Neurobiol 2014; 50: 971–978.

52. Camacho-Navas C, Li L, Poply K et al. Pain assessment using physiological responses/markers in different types of pain: a scoping review. NPJ Digit Med 2026; 9: 64.

