## Supplementary figures for "Hippocampal Neuroinflammation and Altered Peripheral Neurobiological Protein Profile in Experimental Arthritis and Systemic Juvenile Idiopathic Arthritis"

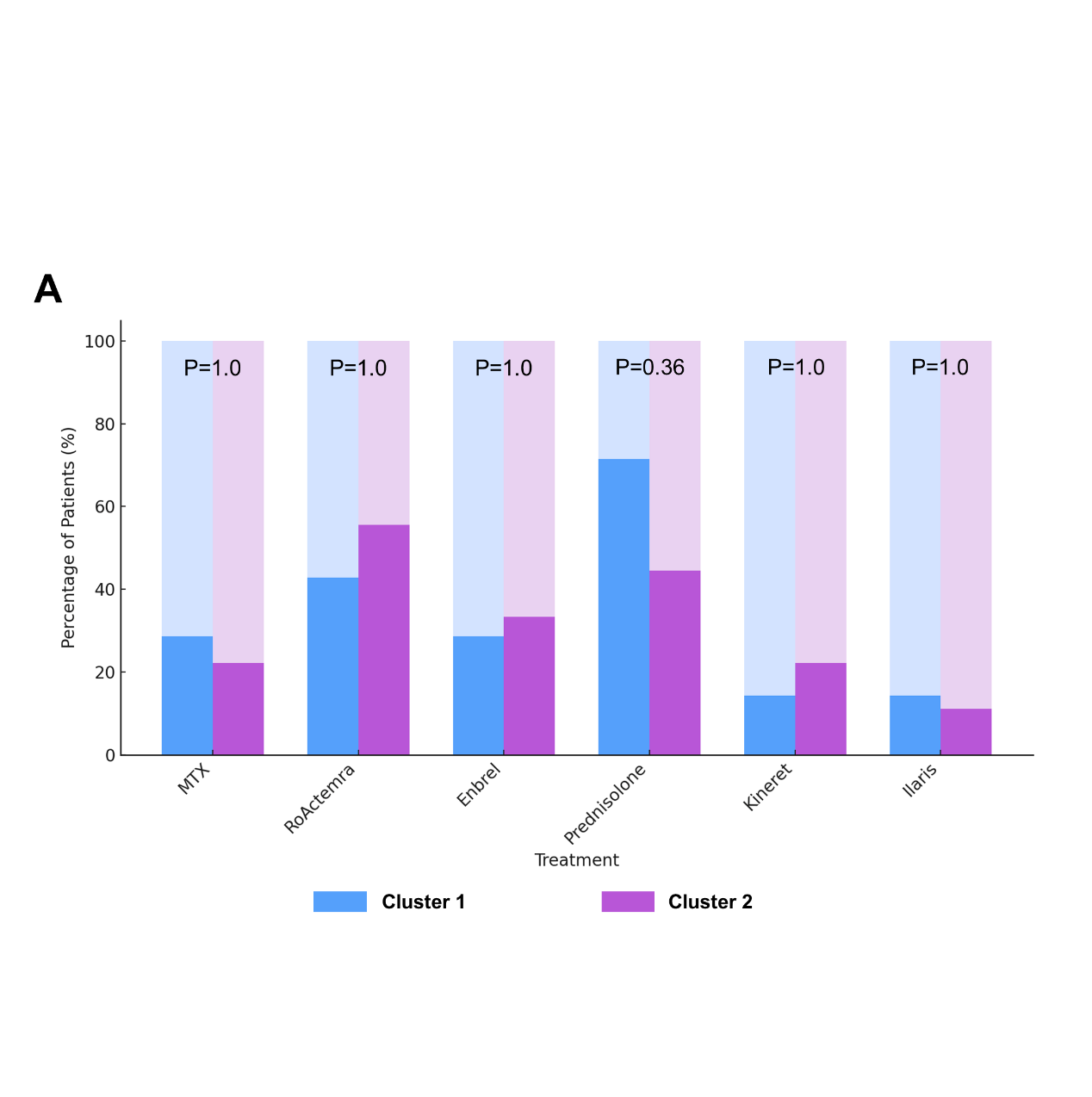

Figure S1. Analysis on the treatment in sJIA patients between two clusters. **(A)** Box plots showing the frequency of different treatments (Enbrel, Ilaris, Kineret, MTX, Prednisolone, and RoActemra) in two clusters of sJIA patients. Statistics: (A) Fisher’s exact test to compare the differences in drug usage frequency between the two clusters. *: *P* < 0.05; **: *P* < 0.01.

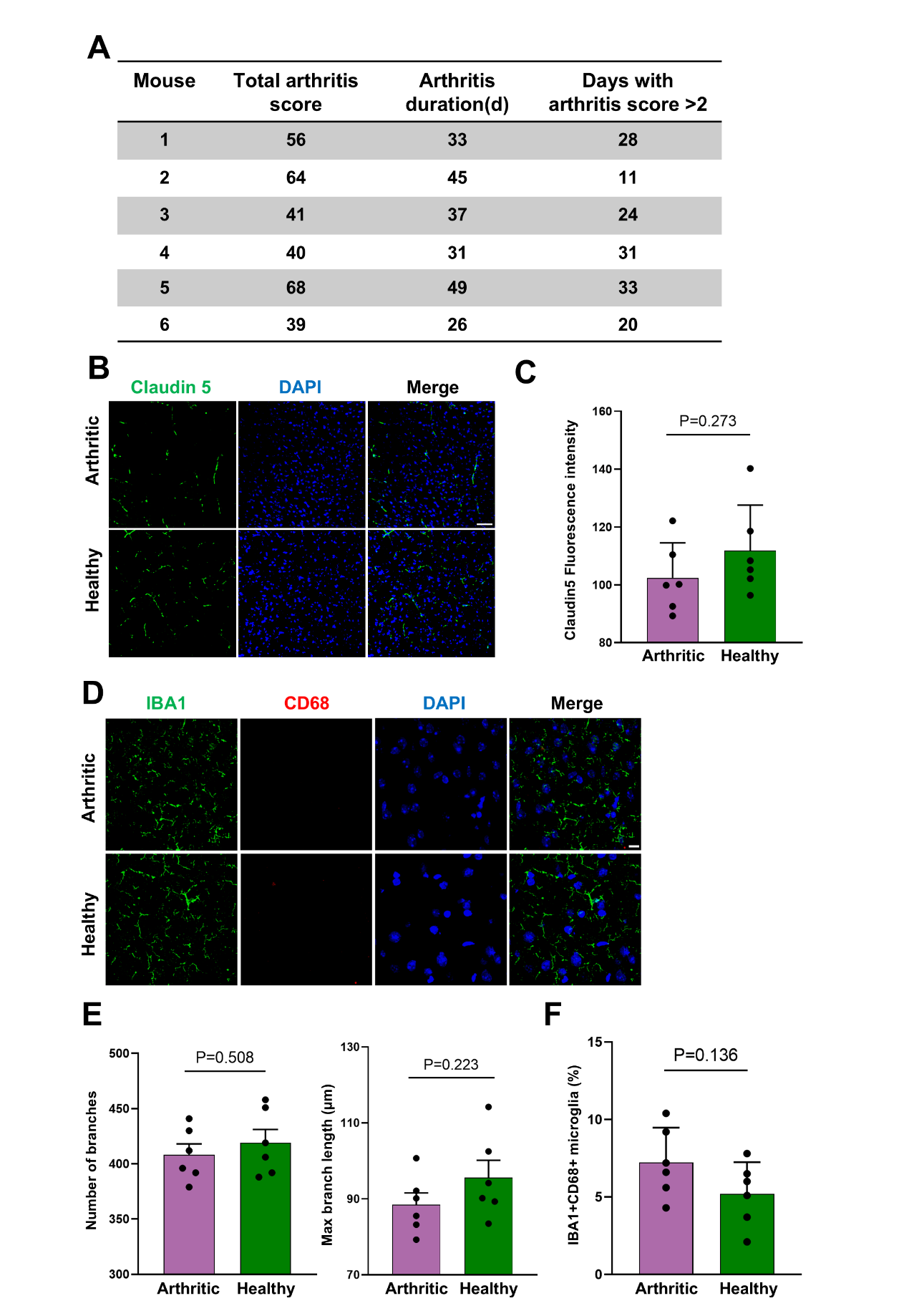

Figure S2. Chronic arthritic mice do not exhibit significant cortical changes compared with healthy mice. **(A)** Detailed arthritis assessment for each mouse (recorded from the booster injection, with 20 time points as referenced in Figure 2A). **(B)** Representative images of IF staining for Claudin-5 in the cortex of arthritic (n = 6) and healthy (n = 6) mice. Scale Bar: 100μm. **(C)** Box plots quantifying Claudin-5 fluorescence intensity by IF in arthritic and healthy mice. **(D)** Representative images of IF staining for microglial markers (IBA1 and CD68) in the cortex of arthritic and healthy mice. Scale Bar: 10μm. **(E)** Box plots quantifying microglial morphology (branch number and branch length) in arthritic and healthy mice. **(F)** Box plots showing the number of activated microglial cells (IBA1⁺ CD68⁺) by IF in arthritic and healthy mice. Statistics: (C, E, F) unpaired t-test.

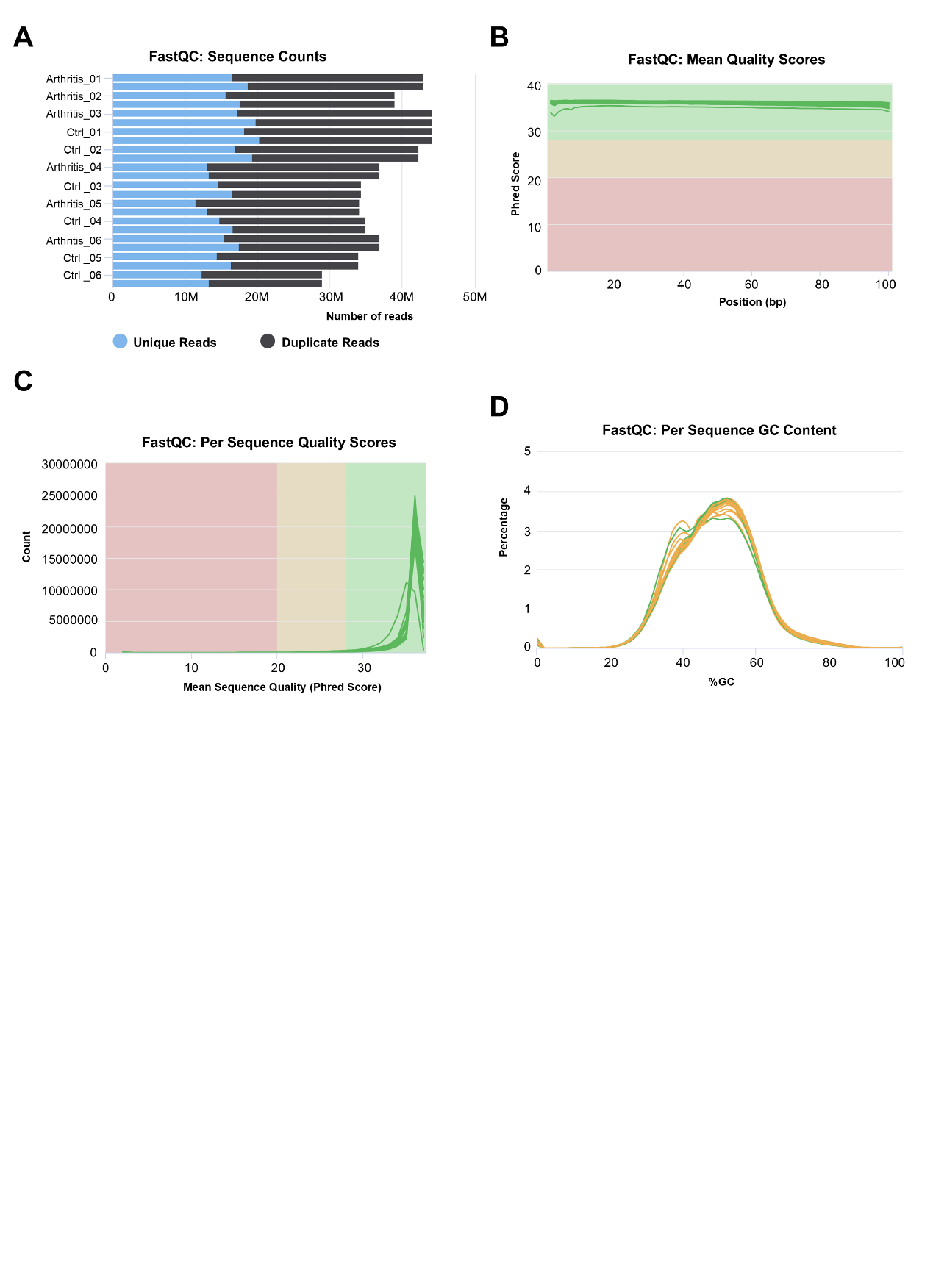

Figure S3. Overview of RNA sequence quality assessed using FastQC and MultiQC. **(A)** Raw sequence counts for each sample, distinguishing between duplicate and unique reads. **(B)** Per-base Phred quality scores, averaged across all bases in each read, indicating high-quality sequencing.

**(C)** Per-sequence Phred quality scores, demonstrating that most reads have a quality >30, confirming their reliability. **(D)** Per-sequence GC content plot, showing a normal distribution of GC content across all sequencing reads.
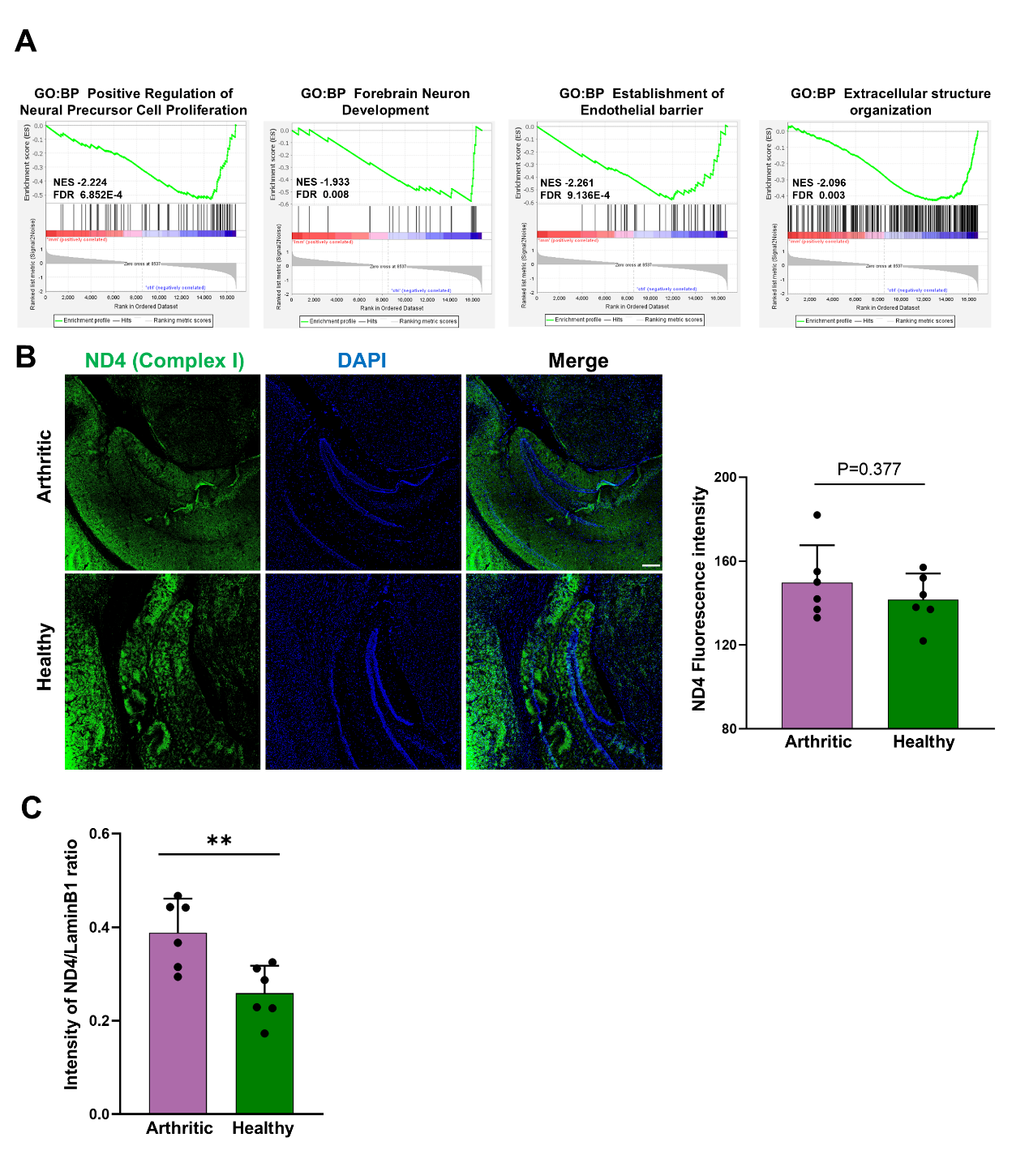

Figure S4. Downregulated pathways and mt-ND4 expression in the hippocampus of arthritic mice compared with Healthy mice. **(A)** GSEA plots showing inhibition of pathways related to neurogenesis and blood–brain barrier integrity in arthritic mice. **(B)** Representative images of IF staining for mt-ND4 in the hippocampus of arthritic (n = 6) and healthy (n = 6) mice, with corresponding box plot quantifying mt-ND4 fluorescence intensity. Scale Bar: 200μm. **(C)** Box plots quantifying mt-ND4 grayscale intensity normalized to Lamin B1 by western blotting (Figure 3E) in arthritic and healthy mice. Statistics: (B, C) unpaired t-test. **: *P* < 0.01.
