## Supplementary tables for "Hippocampal Neuroinflammation and Altered Peripheral Neurobiological Protein Profile in Experimental Arthritis and Systemic Juvenile Idiopathic Arthritis"

**Table S1.** **Demographics and disease characteristics of the study population**

|  | sJIA patients | Healthy controls |
| --- | --- | --- |
| Sample size(n) | 16 | 16 |
| Sex(F) | 8 | 8 |
| Age at sampling (median/range) | 9.0 (3.0-16.0) | 8.0 (4.0-12.0) |
| **CRP** Measurement(n) | 16 |  |
| median/range (mg/L) | 35 (1-193) |  |
| **RF** Measurement(n) | 11 |  |
| positive | 0 (0.00%) |  |
| **ANA** Measurement(n) | 14 |  |
| positive | 2 (14.29%) |  |
| cJADAS-71 | 8.00±2.06* |  |
| VAS pain | 4.39±2.45* |  |
| Life impact | 3.65±2.11* |  |

*: 2 patients lack the scores of these clinical parameters, resulting in 14 patients measured in total.

Abbreviation: C-reactive protein (CRP); Rheumatoid factor (RF); Antinuclear antibodies (ANA); Clinical Juvenile Arthritis Disease Activity Score of 71 points (cJADAS-71); patient visual analogue scale of pain (VAS pain,0-10); patient visual analogue scale of well-being (Life impact,0-10)

**Table S2. Antibodies used in this study**

| Antibodies | Brand and catalog number | Work concentration |
| --- | --- | --- |
| **Primary antibodies (IF)** |  |  |
| Claudin5 | Invitrogen, 34-1600 | 1:250 |
| IBA1 | Abcam, ab5076 | 1:300 |
| CD68 | Abcam, ab125212 | 1:250 |
| DCX | Abcam, ab18723 | 1:300 |
| Mt-ND4 | Invitrogen, PA5-116791 | 1:300 |
| NOX2 | Invitrogen, PA5-79118 | 1:100 |
| 4-HNE | Abcam, ab46545 | 1:200 |
| HMOX2 | Sigma, SAB2501278 | 1:300 |
| P2ry12 | Cell Signaling Technology, 69766 | 1:300 |
| **Primary antibodies (WB)** |  |  |
| IBA1 | Abcam, ab5076 | 1:800 |
| DCX | Abcam, ab18723 | 1:800 |
| Mt-ND4 | Invitrogen, PA5-116791 | 1:1000 |
| HMOX2 | Sigma, SAB2501278 | 1:1000 |
| Lamin B1 | Cell Signaling Technology, 12586 | 1:1500 |
| **Secondary antibodies (IF)** |  |  |
| Alexa Fluor® 488 donkey anti-goat IgG (H+L) | Invitrogen, A-11055 | 1:1000 |
| Alexa Fluor® 647 donkey anti-goat IgG (H+L) | Invitrogen, A-21447 | 1:1000 |
| Alexa Fluor® 647 donkey anti-rabbit IgG (H+L) | Invitrogen, A-31573 | 1:1000 |
| Alexa Fluor® 488 goat anti-rabbit IgG (H+L) | Invitrogen, A-11008 | 1:1000 |
| **Secondary antibodies (WB)** |  |  |
| HRP conjugate-goat anti-rabbit IgG (H + L) | Bio-Rad, 1706515 | 1:2000 |
| HRP conjugate-rabbit anti-goat Immunoglobulins | Dako, P0449 | 1:2000 |

**Table S3. Comparison of neurological protein expression between sJIA and HC in plasma**

| **Protein** | **ΔNPX(sJIA-HC)/Fold Change** | **p_value** | **FDR_BH** |
| --- | --- | --- | --- |
| IMPA1 | -2.27492 | 9.98E-08 | 7.27E-06 |
| ABHD14B | -2.41557 | 2.02E-07 | 7.27E-06 |
| PPP3R1 | -2.09976 | 2.62E-07 | 7.27E-06 |
| FHIT | -1.84467 | 3.20E-07 | 7.27E-06 |
| COL4A3BP | -2.27924 | 4.40E-07 | 8.01E-06 |
| DPEP2 | -1.42284 | 5.41E-07 | 8.21E-06 |
| CRADD | -2.66881 | 7.52E-07 | 9.77E-06 |
| RPS6KB1 | -2.40402 | 1.45E-06 | 1.65E-05 |
| UBE2F | -0.93519 | 2.95E-06 | 2.73E-05 |
| NAA10 | -2.92807 | 3.00E-06 | 2.73E-05 |
| ATP6V1F | -1.90691 | 6.05E-06 | 5.01E-05 |
| HMOX2 | -2.32116 | 7.21E-06 | 5.47E-05 |
| AKT1S1 | -1.37359 | 1.05E-05 | 6.65E-05 |
| KIF1BP | -2.22766 | 1.08E-05 | 6.65E-05 |
| FKBP7 | -1.13078 | 1.10E-05 | 6.65E-05 |
| WWP2 | -1.03245 | 1.21E-05 | 6.65E-05 |
| CETN2 | -1.83528 | 1.24E-05 | 6.65E-05 |
| TBCB | -2.86999 | 1.35E-05 | 6.83E-05 |
| PRTFDC1 | -2.5441 | 2.03E-05 | 9.74E-05 |
| FGFR2 | -0.39734 | 2.66E-05 | 0.000121 |
| ILKAP | -2.05154 | 2.88E-05 | 0.000125 |
| PSME1 | -1.32593 | 7.77E-05 | 0.000322 |
| CD63 | -1.26003 | 0.000110 | 0.000435 |
| GPNMB | -0.19784 | 0.000189 | 0.000715 |
| SMOC1 | 0.827121 | 0.000225 | 0.000819 |
| ECE1 | -1.02307 | 0.000247 | 0.000859 |
| PTPN1 | -2.28161 | 0.000255 | 0.000859 |
| FKBP5 | -2.09816 | 0.000272 | 0.000883 |
| EIF4B | -1.03023 | 0.000355 | 0.001113 |
| ADAM15 | -0.61646 | 0.000517 | 0.001568 |
| DEFB4A | -2.08861 | 0.000651 | 0.001911 |
| PMVK | -1.82161 | 0.000725 | 0.002060 |
| SRP14 | -2.11242 | 0.001040 | 0.002868 |
| EREG | -1.47936 | 0.001120 | 0.002999 |
| PFDN2 | -0.57588 | 0.001393 | 0.003536 |
| CD302 | 0.493304 | 0.001399 | 0.003536 |
| MAD1L1 | -1.97845 | 0.001641 | 0.004004 |
| KIRREL2 | -0.58651 | 0.001672 | 0.004004 |
| LEPR | -0.43325 | 0.001826 | 0.004261 |
| GSTP1 | -0.91604 | 0.002904 | 0.006608 |
| AOC1 | -0.61463 | 0.003183 | 0.007064 |
| ISLR2 | -0.49595 | 0.003407 | 0.007234 |
| IL32 | -0.94637 | 0.003430 | 0.007234 |
| ING1 | -1.8063 | 0.003498 | 0.007234 |
| KLB | -0.77197 | 0.003844 | 0.007774 |
| CDH17 | -0.85533 | 0.003955 | 0.007825 |
| RNF31 | -0.46339 | 0.004933 | 0.009552 |
| BST2 | -0.41492 | 0.005503 | 0.010433 |
| DSG3 | -0.4504 | 0.006241 | 0.011591 |
| TPPP3 | -0.40758 | 0.006665 | 0.012131 |
| NPM1 | -2.29931 | 0.007057 | 0.012592 |
| SFRP1 | -0.69263 | 0.007317 | 0.012804 |
| CTF1 | -0.43732 | 0.008207 | 0.013907 |
| GGT5 | -0.22193 | 0.008252 | 0.013907 |
| ASGR1 | 0.242836 | 0.010252 | 0.016963 |
| PAEP | 1.274969 | 0.010464 | 0.017003 |
| PTS | 0.805303 | 0.012319 | 0.019667 |
| AARSD1 | -0.90518 | 0.016589 | 0.026027 |
| DUSP3 | -0.75734 | 0.022460 | 0.034641 |
| VSTM1 | 0.518674 | 0.029126 | 0.044174 |
| CDH15 | -0.83234 | 0.030238 | 0.045108 |
| NDRG1 | -0.44972 | 0.036639 | 0.053777 |
| IL15 | 0.346412 | 0.041896 | 0.060516 |
| CD33 | 0.533518 | 0.058821 | 0.083637 |
| TNFRSF13C | -0.31789 | 0.071110 | 0.099554 |
| ADGRB3 | -0.13974 | 0.075257 | 0.103763 |
| IL3RA | -0.21028 | 0.084794 | 0.115168 |
| CCL27 | -0.24777 | 0.091647 | 0.122645 |
| RBKS | -0.52251 | 0.110488 | 0.145716 |
| CLSTN1 | -0.23102 | 0.120208 | 0.156270 |
| FCAR | 0.377171 | 0.136439 | 0.174873 |
| PHOSPHO1 | -0.12624 | 0.153056 | 0.193445 |
| PSG1 | 0.345351 | 0.168370 | 0.209886 |
| CEACAM3 | -0.11706 | 0.175144 | 0.215380 |
| PLA2G10 | -0.33581 | 0.226109 | 0.274345 |
| TDGF1 | -0.67336 | 0.274401 | 0.328559 |
| DPEP1 | 0.202431 | 0.302282 | 0.357243 |
| IFI30 | -0.40804 | 0.315414 | 0.367983 |
| IFNL1 | -0.14939 | 0.391039 | 0.450437 |
| NEFL | 0.11106 | 0.407968 | 0.464063 |
| IKZF2 | 0.318182 | 0.452960 | 0.508881 |
| FUT8 | -0.1245 | 0.518434 | 0.575336 |
| HSP90B1 | 0.101921 | 0.533327 | 0.584732 |
| CRIP2 | -0.09166 | 0.585899 | 0.634723 |
| LTBP3 | 0.045228 | 0.649544 | 0.695394 |
| KIR2DL3 | 0.0815 | 0.674287 | 0.713490 |
| ANXA10 | -0.11302 | 0.693869 | 0.725771 |
| SNCG | -0.04925 | 0.727562 | 0.752365 |
| EPHA10 | 0.08658 | 0.873543 | 0.893174 |
| CARHSP1 | -0.00489 | 0.982990 | 0.985944 |
| NXPH1 | 0.003569 | 0.985944 | 0.985944 |

**Table S4. Comparison of neurological protein expression between cluster 1 and cluster 2 sJIA patients**

| **Protein** | **ΔNPX(C2-C1)/Fold Change** | **p_value** | **FDR_BH** |
| --- | --- | --- | --- |
| FKBP5 | -3.15969 | 5.47E-05 | 0.003806 |
| CDH15 | 1.381828 | 8.36E-05 | 0.003806 |
| KIF1BP | -2.15189 | 0.000357 | 0.00778 |
| PRTFDC1 | -2.70749 | 0.00037 | 0.00778 |
| NAA10 | -2.46039 | 0.000427 | 0.00778 |
| PTPN1 | -2.54681 | 0.000696 | 0.008846 |
| TBCB | -2.87358 | 0.000833 | 0.008846 |
| HMOX2 | -2.02248 | 0.000849 | 0.008846 |
| ILKAP | -1.4218 | 0.000875 | 0.008846 |
| CETN2 | -1.47148 | 0.001199 | 0.010913 |
| PMVK | -2.42226 | 0.001459 | 0.011357 |
| COL4A3BP | -1.63856 | 0.001498 | 0.011357 |
| EIF4B | -1.34765 | 0.001847 | 0.012928 |
| KIRREL2 | 0.792989 | 0.002846 | 0.018496 |
| CD63 | -1.45705 | 0.004052 | 0.023703 |
| ASGR1 | -0.31189 | 0.004384 | 0.023703 |
| FGFR2 | 0.322653 | 0.004428 | 0.023703 |
| ECE1 | -0.90944 | 0.006725 | 0.034 |
| DSG3 | 0.68236 | 0.007684 | 0.036716 |
| WWP2 | -1.25308 | 0.008069 | 0.036716 |
| AKT1S1 | -1.48778 | 0.009422 | 0.038497 |
| RPS6KB1 | -1.33026 | 0.009559 | 0.038497 |
| PFDN2 | -0.50155 | 0.00973 | 0.038497 |
| ADGRB3 | 0.345995 | 0.020027 | 0.075937 |
| MAD1L1 | -0.96021 | 0.021399 | 0.077891 |
| CRADD | -1.36491 | 0.025032 | 0.087612 |
| TNFRSF13C | 0.467306 | 0.026643 | 0.089798 |
| SFRP1 | 0.440148 | 0.028641 | 0.093084 |
| FCAR | -0.89771 | 0.034182 | 0.107262 |
| SMOC1 | -0.74864 | 0.036257 | 0.109979 |
| CCL27 | -0.47898 | 0.040185 | 0.117964 |
| VSTM1 | -0.79491 | 0.041677 | 0.11852 |
| FHIT | -0.75313 | 0.047372 | 0.129113 |
| RNF31 | -0.40838 | 0.04824 | 0.129113 |
| IL15 | -0.37235 | 0.052748 | 0.137144 |
| PPP3R1 | -0.84201 | 0.056341 | 0.142417 |
| IFNL1 | 0.464191 | 0.066517 | 0.163597 |
| PSME1 | -0.761 | 0.069897 | 0.167385 |
| FKBP7 | -0.42956 | 0.075118 | 0.175275 |
| DPEP2 | 0.408559 | 0.078134 | 0.177756 |
| SRP14 | -1.00918 | 0.096758 | 0.214756 |
| EREG | -0.534 | 0.119656 | 0.259254 |
| IMPA1 | -0.71611 | 0.133213 | 0.281917 |
| GSTP1 | -0.58532 | 0.138948 | 0.28737 |
| IL32 | 0.577522 | 0.156257 | 0.315986 |
| ATP6V1F | -0.49655 | 0.16342 | 0.316749 |
| CD33 | -0.50494 | 0.163596 | 0.316749 |
| NPM1 | -0.71688 | 0.176532 | 0.334675 |
| ANXA10 | -0.57284 | 0.192596 | 0.354331 |
| AOC1 | -0.43905 | 0.194687 | 0.354331 |
| ING1 | -0.35542 | 0.208023 | 0.367615 |
| CDH17 | 0.546617 | 0.212994 | 0.367615 |
| DEFB4A | -1.03223 | 0.214106 | 0.367615 |
| CTF1 | -0.22727 | 0.240576 | 0.399519 |
| PHOSPHO1 | 0.18474 | 0.245695 | 0.399519 |
| IKZF2 | 0.73201 | 0.245858 | 0.399519 |
| EPHA10 | -0.95448 | 0.265695 | 0.42418 |
| IFI30 | -0.36306 | 0.286625 | 0.442185 |
| PTS | -0.70128 | 0.286692 | 0.442185 |
| NXPH1 | -0.50333 | 0.291838 | 0.442621 |
| NDRG1 | -0.27826 | 0.341255 | 0.50291 |
| CD302 | -0.13903 | 0.342642 | 0.50291 |
| IL3RA | 0.139164 | 0.35064 | 0.50648 |
| BST2 | 0.174796 | 0.366323 | 0.520866 |
| PAEP | 0.791583 | 0.380773 | 0.533082 |
| ABHD14B | -0.32838 | 0.396839 | 0.547156 |
| CARHSP1 | 0.297788 | 0.450408 | 0.611748 |
| GPNMB | -0.04845 | 0.534031 | 0.714659 |
| AARSD1 | -0.25229 | 0.59379 | 0.783114 |
| DUSP3 | -0.08316 | 0.61374 | 0.79661 |
| GGT5 | -0.06265 | 0.632626 | 0.79661 |
| LTBP3 | 0.099321 | 0.634812 | 0.79661 |
| ADAM15 | 0.104781 | 0.639039 | 0.79661 |
| HSP90B1 | 0.107671 | 0.650809 | 0.800319 |
| FUT8 | -0.11978 | 0.667112 | 0.809429 |
| PLA2G10 | -0.14968 | 0.71268 | 0.853341 |
| UBE2F | -0.03576 | 0.735376 | 0.864507 |
| TDGF1 | -0.2575 | 0.750495 | 0.864507 |
| RBKS | -0.15239 | 0.750506 | 0.864507 |
| CEACAM3 | 0.01692 | 0.762485 | 0.867327 |
| NEFL | -0.05875 | 0.794499 | 0.890698 |
| LEPR | 0.046567 | 0.809352 | 0.890698 |
| SNCG | -0.05494 | 0.812395 | 0.890698 |
| PSG1 | 0.086198 | 0.832859 | 0.902264 |
| ISLR2 | 0.055226 | 0.844126 | 0.903712 |
| CRIP2 | 0.033057 | 0.905755 | 0.953429 |
| DPEP1 | 0.032223 | 0.91152 | 0.953429 |
| CLSTN1 | -0.01371 | 0.944907 | 0.977119 |
| KIR2DL3 | -0.01097 | 0.962133 | 0.983754 |
| TPPP3 | 0.007678 | 0.976985 | 0.986244 |
| KLB | 0.00936 | 0.986244 | 0.986244 |

**Table S5. Spearman correlation analysis results between proteins selected and Vas pain score in sJIA patients (ranked by P values)**

| **Protein** | **Spearman_r** | **p_value** | **FDR_BH** |
| --- | --- | --- | --- |
| HMOX2 | -0.679121 | 0.007562 | 0.038862 |
| FKBP5 | -0.670330 | 0.008704 | 0.038862 |
| KIF1BP | -0.670330 | 0.008704 | 0.038862 |
| PRTFDC1 | -0.652747 | 0.011385 | 0.038862 |
| TBCB | -0.648352 | 0.012144 | 0.038862 |
| CETN2 | -0.600037 | 0.023298 | 0.057709 |
| PTPN1 | -0.586813 | 0.027384 | 0.057709 |
| NAA10 | -0.582418 | 0.028855 | 0.057709 |
| COL4A3BP | -0.560440 | 0.037104 | 0.065964 |
| CD63 | -0.538462 | 0.046976 | 0.075162 |
| PMVK | -0.525275 | 0.053748 | 0.076520 |
| EIF4B | -0.512088 | 0.061198 | 0.076520 |
| RPS6KB1 | -0.510451 | 0.062172 | 0.076520 |
| ILKAP | -0.483448 | 0.079885 | 0.091297 |
| WWP2 | -0.463736 | 0.094876 | 0.098468 |
| AKT1S1 | -0.459341 | 0.098468 | 0.098468 |

**Table S6. Spearman correlation analysis results between proteins selected and life impact score in sJIA patients (ranked by P values)**

| **Protein** | **Spearman_r** | **p_value** | **FDR_BH** |
| --- | --- | --- | --- |
| HMOX2 | -0.721673 | 0.003570 | 0.028504 |
| KIF1BP | -0.706271 | 0.004753 | 0.028504 |
| FKBP5 | -0.699670 | 0.005345 | 0.028504 |
| CETN2 | -0.632957 | 0.015115 | 0.050891 |
| TBCB | -0.629263 | 0.015903 | 0.050891 |
| CD63 | -0.607261 | 0.021269 | 0.052839 |
| PRTFDC1 | -0.600660 | 0.023117 | 0.052839 |
| PTPN1 | -0.580858 | 0.029390 | 0.053614 |
| NAA10 | -0.578658 | 0.030158 | 0.053614 |
| COL4A3BP | -0.541254 | 0.045626 | 0.073001 |
| ILKAP | -0.530390 | 0.051042 | 0.074243 |
| WWP2 | -0.514852 | 0.059579 | 0.079438 |
| EIF4B | -0.497250 | 0.070438 | 0.086693 |
| RPS6KB1 | -0.469163 | 0.090569 | 0.100763 |
| PMVK | -0.464247 | 0.094465 | 0.100763 |
| AKT1S1 | -0.389439 | 0.168708 | 0.168708 |

**Table S7. Demographics and disease characteristics of two sJIA clusters**

|  | Cluster 1 | Cluster 2 | P value |
| --- | --- | --- | --- |
| Sample size | 7 | 9 | - |
| Sex(F/M) | 2/5 | 6/3 | 0.315 |
| Age at onset (median/range, year) | 5.2 (2.7-15.0) | 7.4 (0.7-13.8) | 1 |
| Age at sampling (median/range, year) | 5.6 (3.4-16.9) | 9.7 (3.1-15.2) | 0.142 |
| Disease duration (median/range, year) | 0.4 (0.1-2.3) | 4.3(0.5-9.1) | 0.003 |
| **CRP** Measurement(n) | 7 | 9 | - |
| median/range (mg/L) | 21 (1-123) | 36 (1-193) | 0.536 |
| **RF** Measurement(n) | 5 | 6 | - |
| positive | 0 (0.00%) | 0 (0.00%) | - |
| **ANA** Measurement(n) | 6 | 8 | - |
| positive | 2 (33.3%) | 1 (12.5%) | 1 |
| Disease activity score | 7.20±2.37 | 8.60±1.71 | 0.222 |

**Table S8. Top 30 pathways upregulated in arthritic mice (ranked by NES)**

| No. | GO:BP Pathways | ES | NES | FDR q value |
| --- | --- | --- | --- | --- |
| 1 | [AEROBIC_ELECTRON_TRANSPORT_CHAIN](https://www.gsea-msigdb.org/gsea/msigdb/mouse/geneset/GOBP_AEROBIC_ELECTRON_TRANSPORT_CHAIN) | 0.80 | 3.78 | 0.000 |
| 2 | [TRANSLATION_AT_SYNAPSE](https://www.gsea-msigdb.org/gsea/msigdb/mouse/geneset/GOBP_TRANSLATION_AT_SYNAPSE) | 0.90 | 3.55 | 0.000 |
| 3 | [CYTOPLASMIC_TRANSLATION](https://www.gsea-msigdb.org/gsea/msigdb/mouse/geneset/GOBP_CYTOPLASMIC_TRANSLATION) | 0.72 | 3.55 | 0.000 |
| 4 | [MITOCHONDRIAL_TRANSLATION](https://www.gsea-msigdb.org/gsea/msigdb/mouse/geneset/GOBP_MITOCHONDRIAL_TRANSLATION) | 0.64 | 3.49 | 0.000 |
| 5 | [MITOCHONDRIAL_GENE_EXPRESSION](https://www.gsea-msigdb.org/gsea/msigdb/mouse/geneset/GOBP_MITOCHONDRIAL_GENE_EXPRESSION) | 0.62 | 3.32 | 0.000 |
| 6 | [RIBOSOMAL_SMALL_SUBUNIT_BIOGENESIS](https://www.gsea-msigdb.org/gsea/msigdb/mouse/geneset/GOBP_RIBOSOMAL_SMALL_SUBUNIT_BIOGENESIS) | 0.63 | 3.22 | 0.000 |
| 7 | [PROTON_MOTIVE_FORCE_DRIVEN_ATP_SYNTHESIS](https://www.gsea-msigdb.org/gsea/msigdb/mouse/geneset/GOBP_PROTON_MOTIVE_FORCE_DRIVEN_ATP_SYNTHESIS) | 0.80 | 3.20 | 0.000 |
| 8 | [OXIDATIVE_PHOSPHORYLATION](https://www.gsea-msigdb.org/gsea/msigdb/mouse/geneset/GOBP_OXIDATIVE_PHOSPHORYLATION) | 0.67 | 3.19 | 0.000 |
| 9 | [AEROBIC_RESPIRATION](https://www.gsea-msigdb.org/gsea/msigdb/mouse/geneset/GOBP_AEROBIC_RESPIRATION) | 0.61 | 3.18 | 0.000 |
| 10 | [CELLULAR_RESPIRATION](https://www.gsea-msigdb.org/gsea/msigdb/mouse/geneset/GOBP_CELLULAR_RESPIRATION) | 0.54 | 3.10 | 0.000 |
| 11 | [NADH_DEHYDROGENASE_COMPLEX_ASSEMBLY](https://www.gsea-msigdb.org/gsea/msigdb/mouse/geneset/GOBP_NADH_DEHYDROGENASE_COMPLEX_ASSEMBLY) | 0.75 | 3.10 | 0.000 |
| 12 | [MITOCHONDRIAL_RESPIRATORY_CHAIN_COMPLEX_ASSEMBLY](https://www.gsea-msigdb.org/gsea/msigdb/mouse/geneset/GOBP_MITOCHONDRIAL_RESPIRATORY_CHAIN_COMPLEX_ASSEMBLY) | 0.66 | 3.06 | 0.000 |
| 13 | [ELECTRON_TRANSPORT_CHAIN](https://www.gsea-msigdb.org/gsea/msigdb/mouse/geneset/GOBP_ELECTRON_TRANSPORT_CHAIN) | 0.61 | 2.98 | 0.000 |
| 14 | [NUCLEOSIDE_TRIPHOSPHATE_BIOSYNTHETIC_PROCESS](https://www.gsea-msigdb.org/gsea/msigdb/mouse/geneset/GOBP_NUCLEOSIDE_TRIPHOSPHATE_BIOSYNTHETIC_PROCESS) | 0.62 | 2.97 | 0.000 |
| 15 | [RIBOSOME_BIOGENESIS](https://www.gsea-msigdb.org/gsea/msigdb/mouse/geneset/GOBP_RIBOSOME_BIOGENESIS) | 0.54 | 2.93 | 0.000 |
| 16 | [RIBONUCLEOPROTEIN_COMPLEX_BIOGENESIS](https://www.gsea-msigdb.org/gsea/msigdb/mouse/geneset/GOBP_RIBONUCLEOPROTEIN_COMPLEX_BIOGENESIS) | 0.50 | 2.91 | 0.000 |
| 17 | [MITOCHONDRIAL_ELECTRON_TRANSPORT_NADH_TO_UBIQUINONE](https://www.gsea-msigdb.org/gsea/msigdb/mouse/geneset/GOBP_MITOCHONDRIAL_ELECTRON_TRANSPORT_NADH_TO_UBIQUINONE) | 0.83 | 2.90 | 0.000 |
| 18 | [RIBOSOMAL_LARGE_SUBUNIT_BIOGENESIS](https://www.gsea-msigdb.org/gsea/msigdb/mouse/geneset/GOBP_RIBOSOMAL_LARGE_SUBUNIT_BIOGENESIS) | 0.64 | 2.75 | 0.000 |
| 19 | [PROTEIN_TARGETING_TO_MITOCHONDRION](https://www.gsea-msigdb.org/gsea/msigdb/mouse/geneset/GOBP_PROTEIN_TARGETING_TO_MITOCHONDRION) | 0.57 | 2.66 | 0.000 |
| 20 | [MITOCHONDRIAL_RNA_METABOLIC_PROCESS](https://www.gsea-msigdb.org/gsea/msigdb/mouse/geneset/GOBP_MITOCHONDRIAL_RNA_METABOLIC_PROCESS) | 0.61 | 2.66 | 0.000 |
| 21 | [RIBOSOMAL_SMALL_SUBUNIT_ASSEMBLY](https://www.gsea-msigdb.org/gsea/msigdb/mouse/geneset/GOBP_RIBOSOMAL_SMALL_SUBUNIT_ASSEMBLY) | 0.78 | 2.60 | 0.000 |
| 22 | [ENERGY_DERIVATION_BY_OXIDATION_OF_ORGANIC_COMPOUNDS](https://www.gsea-msigdb.org/gsea/msigdb/mouse/geneset/GOBP_ENERGY_DERIVATION_BY_OXIDATION_OF_ORGANIC_COMPOUNDS) | 0.43 | 2.59 | 0.000 |
| 23 | [RIBOSE_PHOSPHATE_BIOSYNTHETIC_PROCESS](https://www.gsea-msigdb.org/gsea/msigdb/mouse/geneset/GOBP_RIBOSE_PHOSPHATE_BIOSYNTHETIC_PROCESS) | 0.47 | 2.57 | 0.000 |
| 24 | [NUCLEOSIDE_TRIPHOSPHATE_METABOLIC_PROCESS](https://www.gsea-msigdb.org/gsea/msigdb/mouse/geneset/GOBP_NUCLEOSIDE_TRIPHOSPHATE_METABOLIC_PROCESS) | 0.50 | 2.56 | 0.000 |
| 25 | [RIBOSOME_ASSEMBLY](https://www.gsea-msigdb.org/gsea/msigdb/mouse/geneset/GOBP_RIBOSOME_ASSEMBLY) | 0.62 | 2.53 | 0.000 |
| 26 | [PROTEIN_RNA_COMPLEX_ORGANIZATION](https://www.gsea-msigdb.org/gsea/msigdb/mouse/geneset/GOBP_PROTEIN_RNA_COMPLEX_ORGANIZATION) | 0.47 | 2.49 | 0.000 |
| 27 | [NUCLEOSIDE_PHOSPHATE_BIOSYNTHETIC_PROCESS](https://www.gsea-msigdb.org/gsea/msigdb/mouse/geneset/GOBP_NUCLEOSIDE_PHOSPHATE_BIOSYNTHETIC_PROCESS) | 0.42 | 2.42 | 0.000 |
| 28 | [PROTEIN_IMPORT_INTO_MITOCHONDRIAL_MATRIX](https://www.gsea-msigdb.org/gsea/msigdb/mouse/geneset/GOBP_PROTEIN_IMPORT_INTO_MITOCHONDRIAL_MATRIX) | 0.76 | 2.41 | 0.000 |
| 29 | [MITOCHONDRION_ORGANIZATION](https://www.gsea-msigdb.org/gsea/msigdb/mouse/geneset/GOBP_MITOCHONDRION_ORGANIZATION) | 0.40 | 2.41 | 0.000 |
| 30 | [MITOCHONDRIAL_TRANSPORT](https://www.gsea-msigdb.org/gsea/msigdb/mouse/geneset/GOBP_MITOCHONDRIAL_TRANSPORT) | 0.47 | 2.40 | 0.000 |

**Table S9. Comparison of inflammation protein expression between sJIA and healthy individuals (ranked by P values)**

| **Protein** | **ΔNPX(sJIA-HC)/Fold Change** | **p_value** | **FDR_BH** |
| --- | --- | --- | --- |
| IL6 | 3.610306 | 2.09E-07 | 1.44E-05 |
| OSM | 1.939349 | 2.24E-05 | 0.000578 |
| TWEAK | -0.60175 | 2.95E-05 | 0.000578 |
| CD5 | -0.85901 | 3.35E-05 | 0.000578 |
| TNFRSF9 | -0.92278 | 4.28E-05 | 0.000591 |
| CD6 | -1.07644 | 0.000102 | 0.001168 |
| MMP-1 | 1.910009 | 0.000579 | 0.00525 |
| SCF | -1.28315 | 0.000609 | 0.00525 |
| EN-RAGE | 2.265121 | 0.000806 | 0.006182 |
| CSF-1 | 0.240953 | 0.001279 | 0.008247 |
| IL-18R1 | 0.588643 | 0.001315 | 0.008247 |
| MMP-10 | -0.65784 | 0.001813 | 0.010423 |
| Flt3L | -0.57402 | 0.00235 | 0.012472 |
| IL18 | 1.970813 | 0.003022 | 0.014893 |
| DNER | -0.30138 | 0.003715 | 0.017088 |
| TGF-alpha | 0.311882 | 0.00456 | 0.019663 |
| CCL23 | 0.629036 | 0.005194 | 0.021081 |
| CXCL6 | -1.25649 | 0.006198 | 0.02376 |
| AXIN1 | -1.08277 | 0.007014 | 0.025473 |
| ST1A1 | -1.1157 | 0.008214 | 0.027137 |
| HGF | 0.530891 | 0.008259 | 0.027137 |
| CDCP1 | 0.485921 | 0.015035 | 0.047156 |
| CASP-8 | -1.02685 | 0.016517 | 0.049551 |
| MCP-1 | 0.471874 | 0.023728 | 0.068217 |
| IL-12B | -0.90411 | 0.027578 | 0.076114 |
| FGF-21 | 1.204073 | 0.031377 | 0.083269 |
| IL7 | -0.71275 | 0.047716 | 0.12194 |
| CD244 | -0.34671 | 0.057056 | 0.140601 |
| CCL19 | -0.53534 | 0.059941 | 0.142618 |
| NT-3 | -0.35086 | 0.066659 | 0.153315 |
| CCL25 | -0.33152 | 0.078907 | 0.175632 |
| IL-17A | 0.542313 | 0.10701 | 0.23074 |
| IL10 | 0.523936 | 0.118612 | 0.248007 |
| MCP-4 | -0.59654 | 0.134077 | 0.272098 |
| uPA | -0.36863 | 0.142213 | 0.280362 |
| CD8A | 0.367393 | 0.159586 | 0.305872 |
| TNFB | -0.51839 | 0.167709 | 0.312755 |
| TRANCE | -0.92792 | 0.190314 | 0.34557 |
| MCP-2 | -0.44523 | 0.231279 | 0.409186 |
| CX3CL1 | -0.23189 | 0.240826 | 0.415424 |
| CXCL5 | -0.56436 | 0.260278 | 0.438029 |
| CD40 | -0.2245 | 0.27533 | 0.452328 |
| STAMBP | -0.49348 | 0.287573 | 0.454841 |
| IL-10RA | 0.352076 | 0.294776 | 0.454841 |
| VEGFA | 0.302714 | 0.296635 | 0.454841 |
| ADA | -0.31895 | 0.305728 | 0.458593 |
| LAP TGF-beta-1 | -0.22785 | 0.404649 | 0.594059 |
| 4E-BP1 | 0.389567 | 0.418868 | 0.602123 |
| CCL3 | 0.226282 | 0.447075 | 0.629555 |
| PD-L1 | 0.128784 | 0.456909 | 0.630535 |
| LIF-R | 0.061156 | 0.514484 | 0.6923 |
| CXCL9 | 0.193756 | 0.521733 | 0.6923 |
| FGF-19 | -0.20881 | 0.54104 | 0.704373 |
| SIRT2 | -0.34184 | 0.552965 | 0.706566 |
| CCL20 | -0.13996 | 0.633569 | 0.794841 |
| IL-10RB | -0.04515 | 0.701675 | 0.864563 |
| TNFSF14 | 0.106461 | 0.751056 | 0.887644 |
| TRAIL | -0.04478 | 0.751955 | 0.887644 |
| FGF-23 | 0.062003 | 0.759 | 0.887644 |
| CCL4 | 0.071418 | 0.805708 | 0.926564 |
| CCL11 | -0.0394 | 0.834 | 0.927593 |
| CXCL1 | -0.08076 | 0.835922 | 0.927593 |
| CCL28 | -0.02496 | 0.858503 | 0.927593 |
| IL8 | -0.04732 | 0.860376 | 0.927593 |
| CST5 | 0.018681 | 0.912796 | 0.939951 |
| IL-15RA | 0.008072 | 0.922829 | 0.939951 |
| CXCL10 | 0.030957 | 0.937208 | 0.939951 |
| CXCL11 | 0.044263 | 0.937365 | 0.939951 |
| OPG | 0.010371 | 0.939951 | 0.939951 |

**Table S10. Comparison of inflammation protein expression between Cluster 1 and Cluster 2 sJIA patients (ranked by P values)**

| **Protein** | **ΔNPX(C2-C1)/Fold Change** | **p_value** | **FDR_BH** |
| --- | --- | --- | --- |
| AXIN1 | -2.471 | 0.000026 | 0.001794 |
| SCF | -1.78449 | 0.000071 | 0.002450 |
| CD40 | 0.830394 | 0.000276 | 0.006348 |
| IL-18R1 | 0.691421 | 0.000595 | 0.010264 |
| TRAIL | -0.55381 | 0.001059 | 0.014614 |
| PD-L1 | 0.669382 | 0.002421 | 0.027842 |
| IL18 | 2.072796 | 0.002980 | 0.029374 |
| VEGFA | 0.917191 | 0.003897 | 0.033612 |
| CCL20 | 0.999251 | 0.004818 | 0.036938 |
| uPA | -0.49929 | 0.005425 | 0.037433 |
| HGF | 0.736415 | 0.006095 | 0.038221 |
| IL6 | 2.394915 | 0.009356 | 0.053797 |
| EN-RAGE | 2.187101 | 0.012916 | 0.068552 |
| IL7 | 0.824604 | 0.015036 | 0.074117 |
| CD6 | -0.4949 | 0.016364 | 0.075274 |
| OSM | 1.13747 | 0.018024 | 0.077730 |
| IL10 | 0.699876 | 0.019209 | 0.077990 |
| CXCL5 | 1.202333 | 0.025734 | 0.098641 |
| IL8 | 0.55269 | 0.029811 | 0.108236 |
| SIRT2 | 1.369262 | 0.031697 | 0.109355 |
| LAP TGF-beta-1 | 0.598075 | 0.034177 | 0.112296 |
| MMP-1 | 1.181673 | 0.038002 | 0.119187 |
| CXCL1 | 1.06217 | 0.042004 | 0.126012 |
| Flt3L | -0.43693 | 0.046144 | 0.132664 |
| CSF-1 | 0.169859 | 0.053551 | 0.147801 |
| LIF-R | 0.205393 | 0.065178 | 0.172978 |
| CXCL11 | 1.136153 | 0.075893 | 0.193949 |
| STAMBP | 0.807651 | 0.084503 | 0.208240 |
| TWEAK | -0.24288 | 0.092132 | 0.218855 |
| CASP-8 | 0.58069 | 0.101131 | 0.232601 |
| FGF-19 | -0.58089 | 0.114143 | 0.254060 |
| TNFB | -0.85174 | 0.127473 | 0.274862 |
| TNFRSF9 | -0.39569 | 0.135019 | 0.282312 |
| CCL23 | 0.451118 | 0.144540 | 0.293337 |
| ST1A1 | 0.612746 | 0.180125 | 0.355108 |
| TNFSF14 | 0.501419 | 0.196993 | 0.377570 |
| IL-12B | -0.39813 | 0.210854 | 0.393214 |
| CX3CL1 | -0.26473 | 0.235151 | 0.426986 |
| IL-17A | -0.48136 | 0.259389 | 0.458919 |
| CCL11 | -0.23045 | 0.268691 | 0.463492 |
| CXCL6 | 0.430719 | 0.279182 | 0.469837 |
| MCP-1 | -0.28882 | 0.291294 | 0.478554 |
| MMP-10 | 0.174663 | 0.313130 | 0.501867 |
| CCL3 | 0.3331 | 0.333277 | 0.522637 |
| TRANCE | 0.318546 | 0.366624 | 0.562158 |
| CD5 | -0.14091 | 0.382973 | 0.574460 |
| OPG | 0.133046 | 0.408998 | 0.599869 |
| 4E-BP1 | -0.4734 | 0.420630 | 0.604656 |
| TGF-alpha | 0.094638 | 0.442046 | 0.622473 |
| MCP-2 | 0.355448 | 0.454862 | 0.627710 |
| CCL25 | -0.12705 | 0.509170 | 0.688877 |
| CCL28 | 0.060386 | 0.542778 | 0.720225 |
| CD8A | -0.15728 | 0.555057 | 0.722943 |
| MCP-4 | 0.228621 | 0.565789 | 0.722943 |
| IL-10RA | 0.205943 | 0.586517 | 0.735814 |
| CXCL10 | -0.26679 | 0.600923 | 0.740422 |
| CXCL9 | -0.14757 | 0.663005 | 0.802585 |
| CCL19 | -0.13994 | 0.674985 | 0.802585 |
| DNER | 0.045284 | 0.693019 | 0.808349 |
| FGF-23 | -0.09791 | 0.704771 | 0.810487 |
| CD244 | 0.032043 | 0.754829 | 0.853284 |
| CST5 | -0.03583 | 0.771785 | 0.858597 |
| FGF-21 | -0.1638 | 0.796575 | 0.872438 |
| IL-10RB | -0.02379 | 0.837826 | 0.903273 |
| CDCP1 | 0.025574 | 0.900920 | 0.950970 |
| NT-3 | 0.011508 | 0.931549 | 0.973893 |
| ADA | -0.01182 | 0.955934 | 0.979528 |
| CCL4 | 0.006458 | 0.974516 | 0.988715 |
| IL-15RA | -0.0015 | 0.991690 | 0.991690 |
